# Phage inducible chromosomal minimalist island (PICMI), a family of satellites of marine virulent phages

**DOI:** 10.1101/2023.07.18.549517

**Authors:** Rubén Barcia-Cruz, David Goudenège, Jorge A. Moura de Sousa, Damien Piel, Martial Marbouty, Eduardo P.C. Rocha, Frédérique Le Roux

**Affiliations:** Sorbonne Université, CNRS, UMR 8227, Integrative Biology of Marine Models, Station Biologique de Roscoff, CS 90074, F-29688 Roscoff cedex, France; Department of Microbiology and Parasitology, CIBUS-Faculty of Biology, Universidade de Santiago de Compostela, Santiago de Compostela, Spain; Ifremer, Unité Physiologie Fonctionnelle des Organismes Marins, ZI de la Pointe du Diable, CS 10070, F-29280 Plouzané, France; Institut Pasteur, Université Paris Cité, CNRS UMR3525, Microbial Evolutionary Genomics, Paris, France; Institut Pasteur, Université Paris Cité, Organization and Dynamics of Viral Genomes Group, CNRS UMR 3525, Paris F-75015 France

**Author notes:** **Corresponding author :** Frédérique Le Roux Equipe Génomique des Vibrios, UMR 8227, Integrative Biology of Marine Models, Station Biologique de Roscoff, CS 90074, F-29688, Roscoff cedex, France.

**Keywords:** vibrio, marine viral particle, mobile genetic element, immune system

## Abstract

Phage satellites are genetic elements that hijack the phage machinery for their own dissemination. However, only few phage satellites have been characterized, and mechanisms by which they influence microbial evolution in nature are unclear. Here we identify a new family of satellites, the Phage Inducible Chromosomal Minimalist Island (PICMI), which is broadly distributed in the marine bacteria *Vibrionaceae*. PICMI is characterized by reduced gene content, does not encode genes for capsid remodeling and packages its DNA as a concatemer. PICMI is integrated in the bacterial host genome at the end of the *fis* regulator and encodes three core proteins necessary for excision and replication. PICMI is dependent on virulent phage particles to spread to other bacteria and confers host protection from other competitive phages, without interfering with its helper phage. The discovery of PICMI strongly suggests that phages, including virulent ones, play important roles for mobility of phage defense elements.

## INTRODUCTION

Bacteriophages (or phages) are viruses that infect bacteria and may be the most diverse and abundant biological entities in the ocean^1, 2^. Since most phages kill their hosts because of their life cycles, phages are key players for promoting bacterial abundance and diversity. Phages are themselves exploited by phage satellites, a class of mobile genetic elements that hijack the phage machinery to promote their own dissemination while interfering with phage reproduction^3–6^. Recently, several studies revealed that marine bacteria transduce a vast diversity of chromosomal islands, many of which might be satellites. These elements were named Virion Encapsidated Integrative Mobile Element (VEIME)^7^ and Tycheposons^8^. These studies suggest that satellites play important roles in natural environments, however their function and characteristics are still poorly understood.

Satellites may develop different strategies for hijacking the life cycle of a helper phage, and the characterization of new satellites can reveal features that unite or separate different satellite families^9^. Much of our knowledge about the lifestyle of phage satellites results from a limited number of elements discovered in clinically relevant bacteria, such as P4-like satellites in *Enterobacterale*s^10^, the phage-inducible chromosomal islands (PICIs) in *Bacillales* and *Gammaproteobacteria*^3,11^, its closely related family capsid forming PICIs (cf-PICI) in Proteobacteria and Firmicutes^12,13^, and the phage-inducible chromosomal islands-like elements (PLEs) in *Vibrio cholerae*^14, 15^. A common feature of all known satellites is their integration in a specific site of the bacterial genome, where satellites remain until their excision is promoted by the induction of a helper prophage, infection by a helper temperate phage, or infection by the virulent helper phage ICP1. This process requires integrases and excision factors encoded by the satellites^6^. The circularized extrachromosomal element then replicates extensively using its own origin of replication and is packaged into viral particles using *pac* and/or *cos* packaging systems^16^. Known satellites typically have genomes about one-third of the size of the helper phages. Many satellites encode diverse mechanisms of capsid remodeling in order to fit their smaller genomes, whilst excluding the larger genomes of their helper phages^6, 10, 15, 17^. Specific features of each phage satellite family include the lifestyle of their helper phage, i.e. temperate for PICI, cf-PICI and P4-like satellites vs. virulent for the helper phage of PLE named ICP1, their genome size of an average 10, 9.5, 14, 18 kbp for P4-like satellites, PICI, cf-PICI and PLE, gene repertoires and the mechanisms they use to subvert the host phage particles. The exploration of known satellites raises the question of the minimal gene set required to excise them from the genome, replicate as an extra-chromosomal element, and hijack the helper phage. Whether the use of its machinery has a cost on helper phage reproduction is also crucial to understand the interaction between the bacterial host, the phage, and the satellite.

The effect of satellites on phage reproduction varies across families, and sometimes even across satellites of the same family. While P4-like satellites and PICIs only interfere partially with the reproduction of their helper phages^18, 19^, PLEs completely abrogate the production of ICP1 progeny^15^. Finally, some P4-like satellites^20^ and PICI^21^ encode hotspots of antiviral systems protecting both the bacterial host and their helper phages from competing phages and other mobile genetic elements. The associations of known phage satellites thus range from pure parasitism to mutualism in relation to their bacterial and phage hosts. While it has become clear that phage satellites are abundant and play diverse roles that affect their hosts, only a limited number have been described, hindering our ability to fully understand the breadth of their influence.

Given the abundance, diversity, and distribution of viruses in the ocean, identifying new phage satellites and understanding their functions might be akin to “looking for a needle in a haystack”. Characterizing and establishing the functions of new phage satellites requires identification of the cognate helper phages and the cellular hosts to understand the parasitic life cycle. In the present study, we addressed this challenge by taking advantage of bacteria from the *Vibrionaceae* family, which is unique for its extensive culture and sequence coverage of hosts and phages^22, 23^. The *Vibrionaceae* (vibrios) comprise a diverse group of bacteria that are widespread within marine environments, encompassing human and animal pathogens^24, 25^. Vibrios are easily cultured, allowing isolation of their infective phages, whole genome sequencing, and inverse genetics.

We report the discovery of a new family of satellites that hijack virulent phages. We named this family of satellites PICMI (<u>P</u>hage <u>I</u>nducible <u>C</u>hromosomal <u>M</u>inimalist Island) because of their reduced size and gene content. When sequencing the genome of a virulent phage and its vibrio host, we detected concatemeric repeat sequences of PICMI in virus particles. PICMI encodes three core proteins necessary for excision and replication but is completely dependent on its helper phage to generate the PICMI infective particles and mobilize across bacteria. Despite this high dependency, PICMI does not strongly interfere with the fitness of its helper phage. However, the satellite can confer host protection from other phages. PICMIs are broadly distributed in the *Vibrionaceae* which encompasses the potential pathogens *V. cholerae*, *V. parahaemolyticus* and *V. vulnificus*^24^, suggesting an important role of this marine phage satellite in *Vibrionaceae*.

## RESULTS

### Identification of a satellite, its cognate host, and helper phage from marine viral particles

We previously isolated and sequenced 49 phages infecting strains of *V. chagasii*^22^. One virulent phage (115_E_34-1, named Φ115 for simplicity) piqued our curiosity because its genome sequence assembly revealed two contigs of 47,851 and 6,110 bp (Fig. S1). The number of sequencing reads was 634,740 and 108,219 for the large and small contig respectively, with only 16 hybrid reads, providing evidence for two distinct mobile genetic elements in the Φ115 phage progeny. The large element corresponds to the genome of the Φ115 phage. The small element contains six genes encoding an integrase (*int*), a putative regulator (*alpA*), a putative primase (*prim*) and three other genes of unknown function (Fig. 1A). Among the genes of unknown function, one gene has four nucleotides overlap (ATGA) with the sequence of *int* and was named *IOLG* for <u>I</u>ntegrase <u>O</u>ver <u>L</u>apping <u>G</u>ene. The two other genes were named *UP1* and *UP2* for <u>U</u>nknown <u>P</u>rotein 1 and 2. Using Phanotate, a tool dedicated to phage genome annotation (see Methods), we identified six additional open reading frames (ORFs), (Fig. S2) which were considered highly questionable because they encoded for proteins of 32 to 44 amino acids. These doubtful ORFs and/or pseudogenes explain the existence of a large non-coding region between *fis* and *UP2*. The 6,110 bp element was also found in the genome of the host used to isolate the phage, *V. chagasii* 34_P_115 (herein named V115), integrated at the end of the *fis* regulator gene and flanked by two direct repeats of 17 bp (Fig. S1). We thus assumed that the 6,110 bp element is a phage satellite and the phage Φ115 is its helper phage.

**Figure 1.**
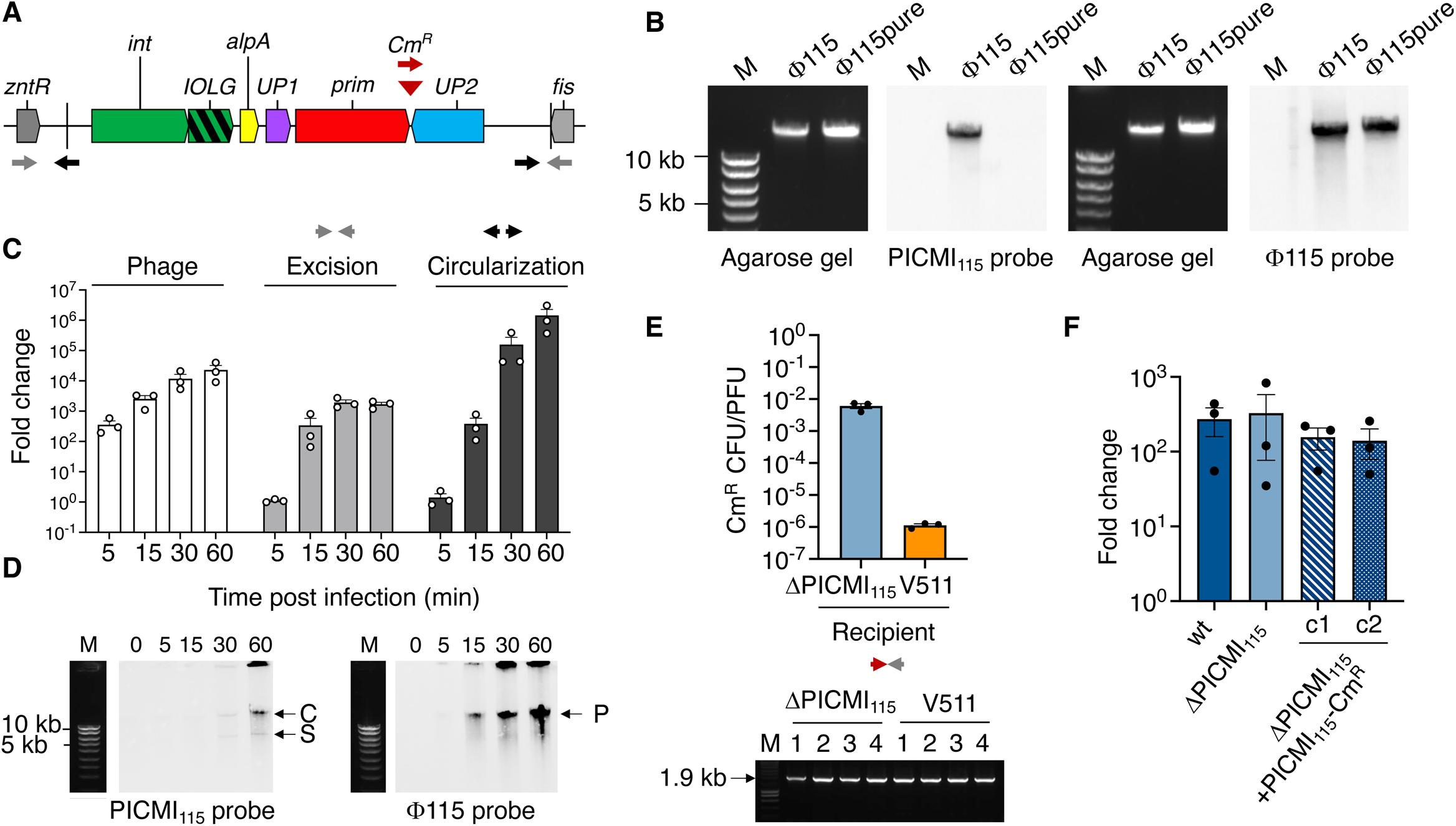
Life cycle of PICMI_115_. **A-**Schematic representation of PICMI_115_ integrated between the *znt*R and *fis* genes. Arrows depict inward- and outward-directed primers used to detect the integration site after excision (grey) or excised/circularized PICMI_115_ (black). For transduction experiments (D) PICMI_115_ was marked by Cm^R^ (brown triangle). Brown forward and grey reverse primers were used to control of the integration of PICMI_115_ at the end of *fis* gene. **B-** Estimation of phage and PICMI DNA size. DNA extracted from phage Φ115 or as control Φ115pure, were separated on an agarose gel (left panel, SYBR Green stained), and Southern blotted (right panel) with PICMI_115_ or Φ115 probes. M: molecular marker (Smart ladder Eugentec). **C** and **D-**The dynamics of excision and replication were explored by qPCR (C) and Southern blot (D). Bar charts show the mean fold change +/- SEM. from three independent experiments (individual dots). In the southern blot C, S and P indicate the concatemeric and single form of PICMI_115_ and phage genome respectively. **E-** To determine transduction frequencies, the phage Φ115pure was produced from a derivative of V115 carrying a Cm^R^ marked PICMI_115_ (see A). ΔPICMI_115_ and V511were used as recipient cells. Results (upper panel) indicate the ratio between the titer of PICMI_115_-Cm^R^ (CFU/ml) and the titrer of phage (PFU/ml). Bar charts show the mean +/- SEM from three independent experiments (individual dots). The integration of PICMI_115_-Cm^R^ at the end of the *fis* gene was confirmed by PCR (SYBR Green gel stained, lower panel). **F-** Interference with the reproduction of the helper phage. The strain V115 wild type, a derivative lacking the full PICMI_115_ and two clones (c1 and c2) of transductants carrying PICMI_115_-Cm^R^, were infected by Φ115pure at MOI10 for 60 minutes. Bar charts show the mean fold change of phage titer +/- s.d. from three independent experiments (individual dots). Differences between treatments are not statistically significant (ANOVA, P=0.75, F-test).

Despite the small size of the satellite (∼1/8 the phage genome), we did not observe smaller-sized capsids commensurate to its genome size, as described for other known families of phage satellites (Fig. S3). We tested for the presence of physical contacts between the two genomes^26^ to confirm that the satellite was located in viral particles lacking the phage genome. We applied HiC^27^ on different mixes of phage particles (see Methods). The result showed a clear absence of physical contact between the DNAs of the phage Φ115 and the satellite (Fig. S4), confirming the exclusive packaging of the satellite in full-sized phage-like particles. To understand if the satellite fills the capsid by packaging as a concatemer, we performed single-molecule nanopore sequencing of Φ115- encapsidated high–molecular weight DNA and found a fraction of the viral particles contained a concatemer of 8 copies of the satellite of ∼49 kbp size, similar to the Φ115 genome (Fig. S5, Table S1). Finally, we confirmed by Southern blot that concatemeric repeat sequences of the satellite show DNA of similar size to that of the genome of phage Φ115 packaged in viral particles (Fig. 1B). To estimate the percentage of Φ115 particles that contain the satellites instead of phage DNA, we first normalized the Illumina sequencing reads on genome size (634,740/47,851=13.26 for the phage and 108,219/6,110=17.71 for the satellite) and next considered that eight copies of the satellite are packaged as concatemer in particles (17.71/8=2.21). This led to 16% (2.21/13.26*100) of the population being hitchhiked by the satellite. This estimation was further confirmed by nanopore sequencing (10 or 15% depending on the replicate, Table S1) and qPCR of phage DNA (15%, Fig. S6). Altogether, these data strongly suggest the identification of a new marine phage satellite. Due to its reduced size, we named this satellite PICMI_115_, for Phage Inducible Chromosomal Minimalist Island identified in vibrio V115 and its cognate helper phage Φ115.

### PICMI sustains minimal function of excision and replication

Our cultivation-enabled model system has enabled us to dissect the various steps in the PICMI’s life cycle: (i) excision, (ii) replication, (iii) packaging (iv) transduction to a new host. Of these, we were not able to identify *cos* or *pac* packaging sites in the helper phage genome, or any homologs of genes involved in redirecting packaging that are characteristic of other satellite families (i.e., *terS*, *sid*, *ppi*). However, Nanopore sequencing of the viral particles revealed random extremities of the PICMI_115_ concatemer (Fig. S5), which is indicative of a mechanism of headful (*pac*-like) packaging.

Prior to exploring the induction of PICMI_115_ by the helper phage, we generated Φ115 viral particles without the PICMI_115_, by two passages in a V115 derivative lacking the entire PICMI_115_. The vibrio mutant was named ΔPICMI_115_, and the phage progeny was named Φ115pure. The absence of the satellite in the Φ115pure population was confirmed by nanopore sequencing (Table S1), qPCR (Fig. S6), and Southern blot (Fig. 1B). To test for helper phage dependent excision/circularization activation, we performed qPCR analysis with inward- and outward-directed primers as shown in Fig.1A. Amplicons were obtained 15 minutes (min) after adding the phage to the bacterial culture (Fig. 1C), the estimated time for its complete adsorption and phage DNA injection in the host cytoplasm (Fig. S7). Excision/circularization was observed exclusively in the presence of the helper phage (Fig. S8). Outward-directed primers also amplify the junctions in the concatemer. A dramatic increase of the copy number of the circular DNA was observed 30 min after phage addition (Fig. 1C), indicating intensive replication. The increase in the amount of a DNA band at the size of the concatemer is observed by Southern blot at 60 min (Fig. 1D), consistent with a plasmid-like rolling circle DNA replication mechanism.

It is expected that PICMI_115_ transduction requires the helper phage to adsorb on the recipient host. The phage Φ115 was previously described as having a narrow host range^22^, with only 2 out of 136 *V. chagasii* strains (V115 and V157) susceptible to Φ115 infection and reproduction. These two strains each encode one (identical) PICMI. Testing closer phylogenetic neighbors to the original host V115, we found that the phage Φ115 adsorbs to the strain V511 without producing progeny (Fig. S9), probably due to intracellular defense mechanisms^23, 28^. V511 does not carry PICMI and shows 100% identity with the *fis* gene of V115 and V157. We thus assumed that the vibrio strain V511, in addition to the V115 derivative lacking PICMI_115_ (ΔPICMI_115_), could be used as recipient for transduction assays. We first inserted a chloramphenicol resistance marker (Cm^R^) downstream of the *prim* gene of PICMI_115_ (Fig. 1A) and infected this strain with Φ115pure to produce lysates of viral particles with the Cm^R^- encoding PICMI_115_. The introduction of the Cm^R^ cassette slightly increased the copies number of PICMI_115_ in phage particles (Fig. S6). We thus assumed that the excision, replication, and packaging functions of the PICMI_115_-Cm^R^ satellite were intact. We next used this lysate to infect the two susceptible hosts, ΔPICMI_115_ and V511, and selected for chloramphenicol resistant cells that acquired the PICMI_115_-Cm^R^ satellite. Transductants were obtained at a multiplicity of infection (MOI) from 0.1 to 0.0001 for the recipient ΔPICMI_115_, and MOI from 10 to 0.01 for V511. PICMI_115_-Cm^R^ was transduced at higher frequency when ΔPICMI_115_ was the recipient (transductants in CFU.ml^-^1/ phages in PFU.ml^-^1∼ 6.10^-^3) relative to V511 (10^-^6) (Fig. 1E, upper panel). We confirmed by PCR that the integration of PICMI_115_-Cm^R^ occurred at the end of the *fis* gene in all tested transductants (Fig. 1E, lower panel). Altogether our results showed that PICMI_115_ is activated by a virulent phage, replicated by rolling circle, packaged, and transduced as a concatemer. With roughly 15% of viral particles that contain the satellite, the question arises whether PICMI_115_ inflicts a cost on its helper phage.

Most known satellites interfere at least partially with the reproduction of their helper phage, although this effect can vary broadly between families^15, 18, 19^. We thus quantified the extent to which PICMI115 interferes with its helper phage Φ115. We compared the titer of phages produced by the bacterial strains V115 wild type (wt), ΔPICMI_115_, and two clones of ΔPICMI_115_ +PICMI_115_-Cm^R^ after infection by Φ115pure. No significant differences (ANOVA, P=0.75, F-test) were observed between the strains (Fig. 1F), showing that PICMI_115_ does not strongly impact the reproductive fitness of its helper phage.

### AlpA is a key regulator of PICMI activation

Having established that PICMI_115_ is a phage satellite, we further analyzed the role of each PICMI_115_-encoded gene in the various aspects of the satellite’s lifestyle. Each of the six genes (Fig. 1A) was deleted in V115, and the mutants were compared to the wild-type host for excision, packaging, and transduction. This revealed that *alpA, int,* and *IOLG* are necessary for PICMI_115_ excision induced by the helper phage (Fig. 2A; Fig. S10). The deletion of *prim* results in a lower number of circularized PICMI_115_ copies, relative to the wild type, indicating that the primase is involved in PICMI_115_ replication (Fig. 2A; Fig. S10). Accordingly, the number of viral particles that contain PICMI_115_ in phage progenies was strongly reduced in Δ*alpA,* Δ*int*, Δ*IOLG,* and Δ*prim* deletion mutants (Fig. 2B, Fig. S6). It also led to a much lower transduction of PICMI_115_-Cm^R^, below our limit of detection (Table 1). When expressed in trans in the V115 derivative Δ*int*, *int,* with or without the overlapping gene *IOLG,* restored the helper phage-induced excision of PICMI_115_ (Fig. 2C). The expression of *int/IOLG* also complemented the Δ*IOLG* deletion (Fig. 2C). Notably, the expression of *alpA* in trans was sufficient to induce PICMI_115_ excision even in the absence of phage (Fig.2C), and the copy number of circularized satellites slightly increased 30 min post infection (Fig. S11).

**Figure 2.**
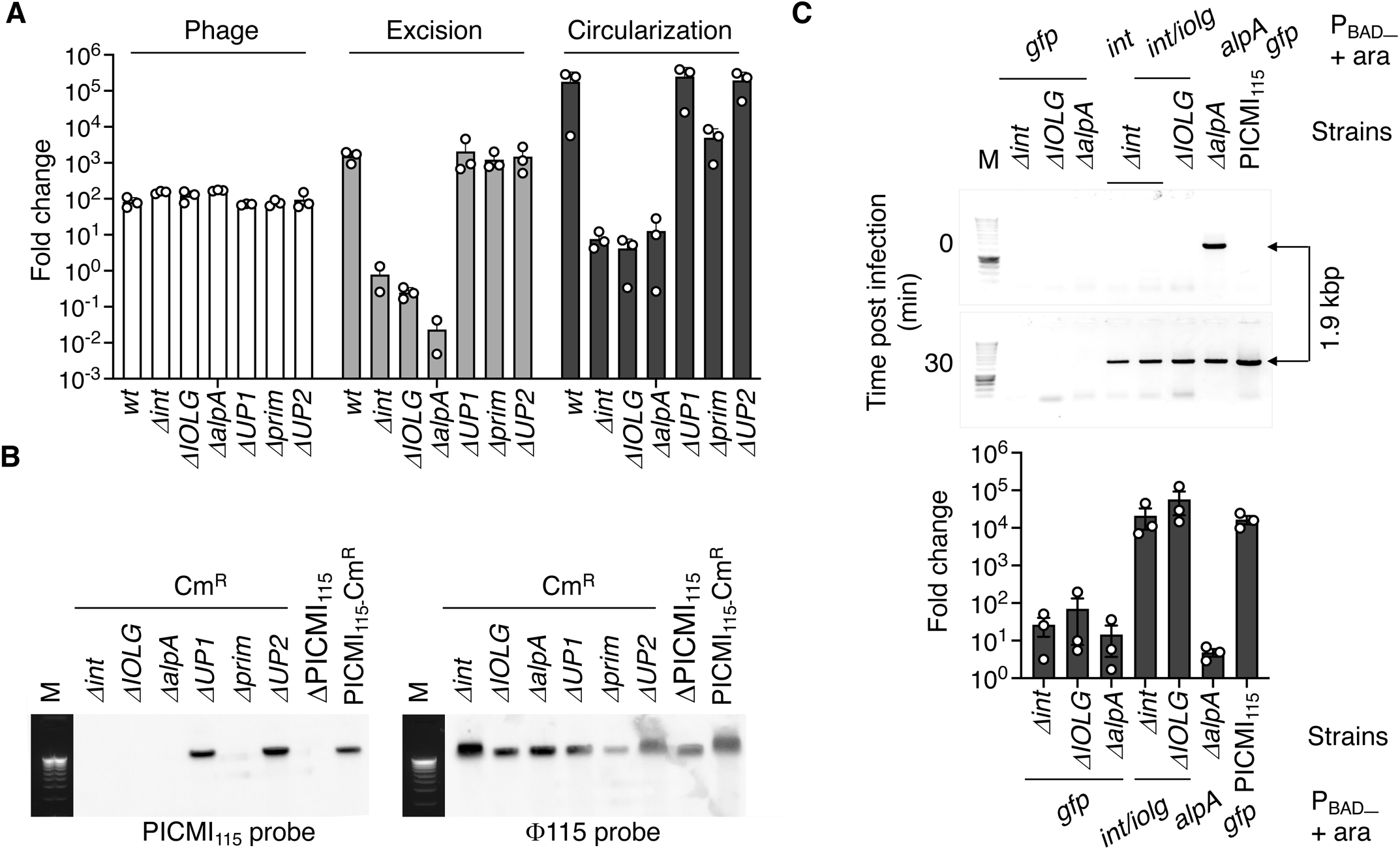
Genes involved in PICMI_115_ activation. **A-** Fold change of the copies number for phage, empty integration site and circularized PICMI_115_, 30 minutes post-infection by the phage F115pure. Bar charts show the mean +/- SEM. from three independent experiments (individual dots). **B-** Phage progenies were produced using DPICMI_115_, PICMI_115_-Cm^R^ and derivatives lacking a single gene (e.g. Dint). Phage DNAs were separated on an agarose gel and Southern blotted with PICMI_115_ or F115 probes. M: molecular marker (Smart ladder Eugentec). These phages were also used for transduction experiments (Table 1). **C-** For complementation assays, the genes necessary for PICMI_115_ excision, *int/IOLG* and *alpA* or, as control, *gfp* were cloned under the control of the arabinose inducible. Circular from of PICMI_115_ was detected by classical PCR and gel stained (upper panel) or qPCR (fold change 30 min post infection, lower panel). Bar charts show the mean +/- SEM. from three independent experiments (individual dots). The expression of *alpA* is sufficient to induce DPICMI_115_ activation in the absence of phage (upper panel, time 0 min), explaining the similar fold change between the *Dalp* mutant and its complemented derivative (lower panel).

**Table 1.** PICMI_115_ transfer by Φ115 phage produced from different donors

| **Donor strain** | **PICMI<sub>115</sub> titre<sup>a</sup>** |
| --- | --- |
| PICMI<sub>115</sub>-Cm<sup>R</sup> | 5,85E+03 ± 1,62E+03 |
| Δ*int*-Cm<sup>R</sup> | <1 |
| Δ*IOLG*-Cm<sup>R</sup> | <1 |
| Δ*alpA*-Cm<sup>R</sup> | <1 |
| Δ*UP1*-Cm<sup>R</sup> | 4,78E+03 ± 1,98E+03 |
| Δ*prim*-Cm<sup>R</sup> | <1 |
| Δ*UP2*-Cm<sup>R</sup> | 4,88E+03 ± 1,52E+03 |

<sup>a</sup>PICMI<sub>115</sub> titer/ml of lysate, using vibrio ΔPICMI<sub>115</sub> as recipient strain. The titer of phages produced from the different donors was 10<sup>6</sup> PFU/ml. The means and standard deviations from four independent experiments are presented.

AlpA is predicted to act as a DNA-binding regulator, and previous work suggested that it is a transcriptional regulator^3^ and/or an excisionase^29^. In PICMI_115_, the expression of *alpA* from a plasmid did not alter the expression of the satellites’ genes, suggesting that *alpA* induction of PICMI_115_ excision is not mediated by transcriptional regulation (Fig. S12A, B). We used ColabFold^30^ to search for structural similarities with known excisionases and found strong similarities with the Torl regulator in *E. coli* and Xis excisionase from *Streptomyces ambofaciens* (Fig. S12C). We conclude that among the three genes essential for excision, *int* and *IOLG* are constitutively expressed, *alpA* is activated by phage, and the three proteins are involved in the formation of the excision complex.

### The intensity of PICMI_115_ activation depends on the phage used as helper

The induction of the satellite can be highly specific to the helper phage(s)^5^. We searched for other phages that could infect the strain V115 to establish whether induction of PICMI_115_ is also specific to the helper phages Φ115. We used viruses from seawater collected at the same oyster farm four years later (see Methods) and isolated seven phages infecting the host V115. The probe directed against phage Φ115 also detected the newly isolated phages by Southern blot, suggesting that they are genetically related (Fig. 3A). Except for Φ27, PICMI_115_ was detected in the progenies resulting from the infection of all other phages, although at a much lower quantity than in the Φ115pure infection (Fig. 3A). The phage Φ27 was unable to induce a detectable excision of PICMI_115_ (Fig. 3B). We thus compared the expression of the PICMI genes upon Φ115pure (Fig. 3C) and Φ27 infection (Fig. 3D). This revealed that 15 min post infection by Φ115pure, presumably as soon as the phage is injected in the cytoplasm (Fig. S7), only *alpA*, *UP1* and *prim* were upregulated, and thus defined as an early regulon. An increase of transcripts for the remaining PICMI_115_ genes (*int, IOLG* and *UP2*) was observed after 60 min. This resulted from the increase of the number of copies of the PICMI_115_ genome and not necessarily through gene activation. Infection by phage Φ27, which did not lead to PICMI_115_ induction, has no reproductible effect on PICMI_115_ gene expression (Fig. 3D). Altogether our results demonstrate that Φ115 spurs strong induction of PICMI_115_ and that the ability to induce PICMI is related to the ability to efficiently induce the early PICMI_115_ regulon.

**Figure 3.**
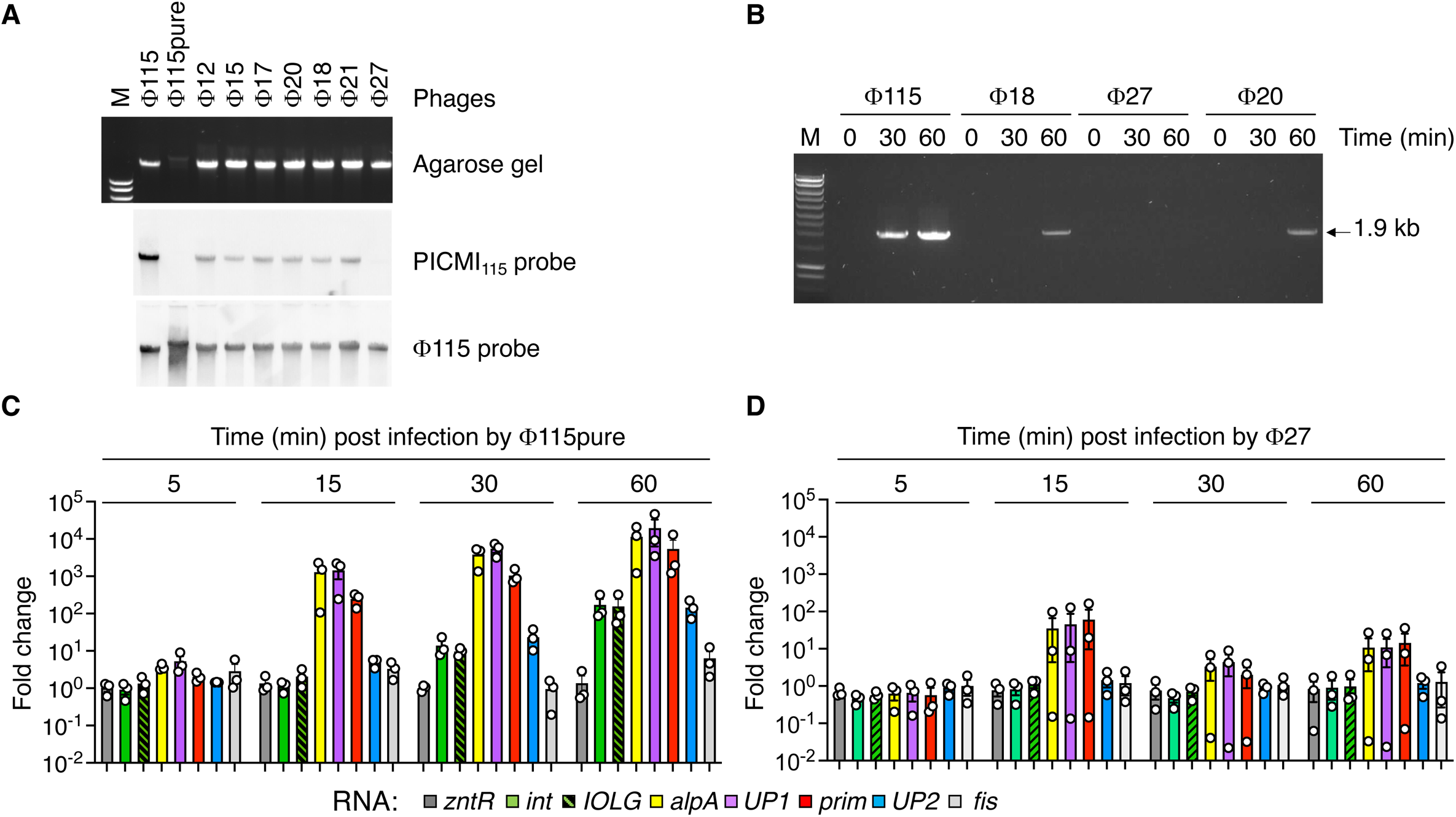
The activation of PICMI_115_ is helper phage specific. **A-** PICMI_115_ DNA was less or not detected in F115 genetically related phages. DNA from viral particles were extracted and separated on SYBR Green stained gel and Southern blotted with an PICM_I115_ or F115 probes. M: molecular marker (Smart ladder Eugentec). **B-** An efficient induction of the satellite is specific to the helper phage F115. The vibrio V115 carrying a Cm^R^ marked PICMI_115_ was infected with the diverse phages at a MOI of 10 for the indicated time, PCR amplicon corresponding to the circularized and concatemeric form of PICMI_115_ were visualized on agarose gel. **C** and **D-** The vibrio V115 was infected with F115pure (**C**) or F27 (**D**). Each of the six genes from PICMI_115_, the two flanking genes *fis*, *zntR* and the house keeping gene *gyrA* were detected by qRT-PCR. Bar charts (same color code than in Fig. 1A) show the mean +/- SEM fold change from three independent experiments (individual dots).

### PICMI-like elements are broadly distributed in the *Vibrionaceae*

Our description of the PICMI minimalism was based on a single model system identified in our collection, raising questions about the size, distribution, and diversity of PICMI-like elements in bacterial genomes. We thus built a MacSyFinder model^31^ to allow the automatic identification of this element in bacterial genomes. Our PICMI_115_ prototype is integrated in the V115 host genome at the end of the *fis* regulator gene and contains only six genes. Among those, we showed that *int, alpA,* and *prim* are necessary for PICMI_115_ lifestyle. In accordance with the experimental data, we set the presence and colocalization of *int, alpA, prim,* and *fis* genes as mandatory in the model. We then used it to search for PICMI-like elements in all Genbank bacterial complete genomes (v243, 05/26/2021) and identified 135 elements (Table S2). From this list, the 67 satellites in *Vibrionaceae* genomes have a significantly smaller size (average 6.7 kbp, unpaired t test P<0.0001) (Fig. S13). We thus extended our search for such elements to a much larger albeit specific dataset of 19185 *Vibrionaceae* genomes (NCBI Assembly database, 02/16/2023). We identified a total of 97 elements, broadly distributed in diverse *Vibrionaceae* species (Fig. 4, Table S3). We never detected more than one PICMI-like element in these genomes, contrasting with P4-like satellites in *E. coli* genomes, that can contain up to three P4-like satellites^10^. Pairwise alignment of the DNA sequences of PICMI- like satellites permitted grouping them into 35 distinct PICMI subfamilies (>90% global pairwise nucleotide identity) (Fig. S14, Table S3). Up to seven subfamilies could be detected in one species (*V. cholerae*) (Fig. S14). In most cases, the distribution of PICMI subfamilies coincides with the host species phylogeny. This is expected for mobile genetic elements transduced by vibriophages that, for the vast majority, have a narrow host range^22, 23, 32^. However, PICMI_115_ was detected in two strains of *V. chagasii* (V115 and V157) isolated during the same time series sampling in France^22^ and in a *V. toranzoniae* strain isolated from cultured clams in NW Spain (cmf 13.9) (Fig. 4 and Fig. S15). Another *V. toranzoniae* isolate from seawater in SE Spain (96-373) carries a different PICMI subfamily. This incongruence between the subfamilies of PICMI and the bacterial hosts suggests that this satellite might be horizontally transferred between diverse *Vibrio* species.

**Figure 4.**
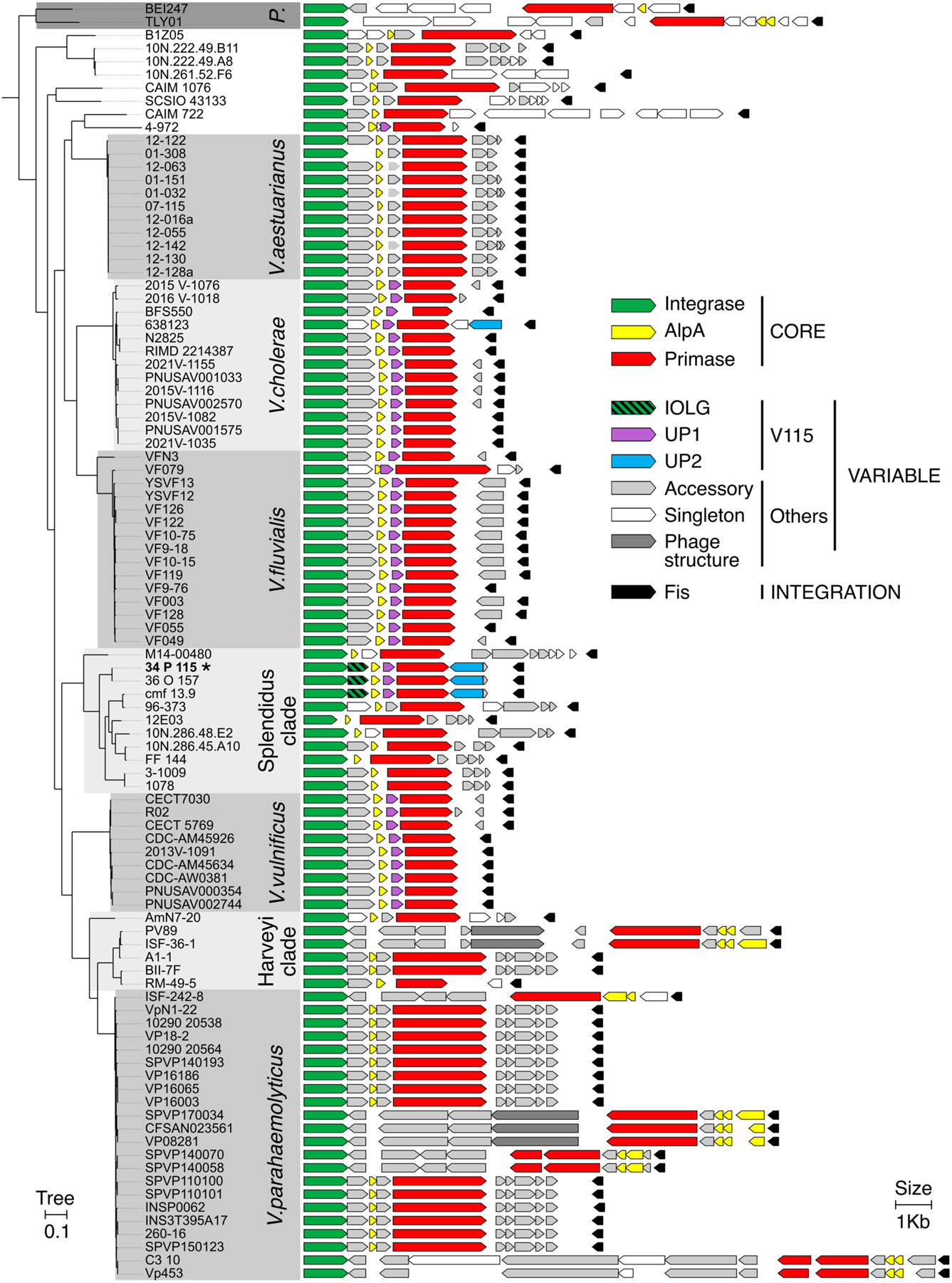
PICMI-like satellite distribution and gene content in the *Vibrionaceae*. Phylogenetic persistent core tree and genomic representation of the 97 PICMI elements found in *Vibrionaceae* (GenBank 01-27-23 containing 19189 organisms). Genus, super clades or of species names are indicated in the grey boxes. “P.” correspond to *Photobacterium* genus, Harveyi, Splendidus are super clades encompassing several *Vibrio* species. The PICMI_115_ element is pinpointed by bold strain name (34_P_115) and by an asterisk. Solid colors indicate core PICMI_115_ genes. Grey colors indicate accessory and singleton PICMI-like genes defined using reciprocal best-hit with 20% identity for 50% coverage.

We then analyzed the genes that are frequent across the PICMI variants. As for PICMI_115_, large non-coding regions might be explained by the presence of false ORF and/or pseudogenes (Fig. S16). Since these genes are not predicted to be functional, they were not considered in our analysis. The core genes encoding the integrase, the AlpA regulator and the primase were identified in all the elements, as expected, as they were used to identify them. We showed above that a gene encoding unknown function overlapping *int* on four nucleotides (*IOLG*) is essential for the excision of the PICMI_115_. Among the 35 PICMI subfamilies, 21 carry a *IOLG* homolog that overlaps *int* over 1, 4 or 13 nucleotides, ATGA being the most frequent overlapping site (n=14) (Fig. 4 and Fig. S14). Four elements carry a gene contiguous to the *int* gene (ICG) and three do not carry a gene between *int* and *alpA*. The *IOLG or ICG* were grouped in 13 distinct gene families (>20% protein identities, 80% LmaxRap), all of them of unknown function. The remaining seven PICMI elements were more divergent in gene repertoires and gene order. In six of them, *alpA* was in two or three copies. These loci may thus have been the result of multiple events of integration, gene loss, and recombination with other satellites. *UP1* homologs were found in 12 subfamilies of PICMI (Fig. 4 and Fig. S14), in line with our observations that *alpA*, *UP1,* and *prim* form the early regulon activated by the helper phage (Fig. 3). In 26 other PICMI subfamilies, single genes were also present between *alpA* and *prim*, forming eight distinct families, each encoding an unknown function. On the nine remaining subfamilies, *alpA* was adjacent to *prim*. Altogether, our analysis revealed that PICMI-like elements have small genomes, are widely distributed in the *Vibrionaceae,* and encode a limited number of genes that are essential for its lifestyle.

### Identification of a new defense system in PICMI_115_

In spite of the small size of PICMIs, all subfamilies have accessory genes and some of them encode for known phage defense systems (Table S3), namely Restriction modification systems type I and II, a retron type II, and Paris type I^33^. These systems are located in the locus between *prim* and *fis,* suggesting that this might be a hotspot for the acquisition of anti-viral defense genes, akin to the locus between the integrase and Psu in P4-like satellites^20^. This also suggests that PICMIs can provide viral defense mechanisms to their host, as observed both in P4-like satellites and in PICI^21^. Consistent with this, we observed that the presence of PICMI_115_ in V511 (transductants) greatly affected the infection outcome of this bacteria by the phage Φ511(Fig. 5A), showing an antiviral effect of the PICMI115.

**Figure 5.**
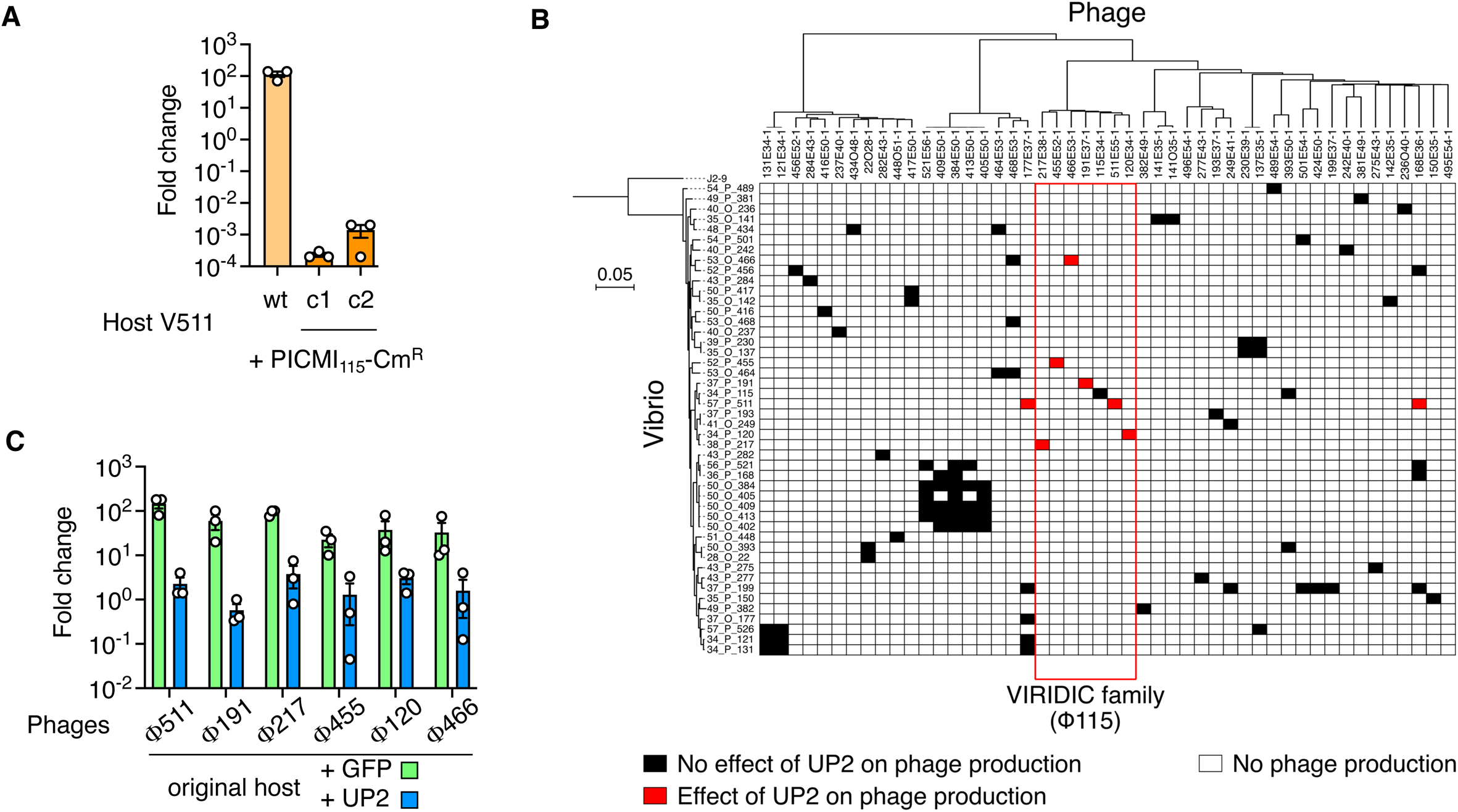
PICMI_115_ and UP2 confer host immunity to specific phages. **A-** PICMI_115_ greatly affected the infection outcome by the phage Φ511. Vibrio strain V511 (wt) and transductants carrying the full satellite (PICMI_115_-Cm^R^, two clones c1 and c2) were infected with F511 at an MOI 10 for 60 minutes. Bar charts show the mean fold change of phage titer +/- SEM from three independent experiments (individual dots). **B-** UP2 encodes a novel defense system which activity depends on the host genetic background. A plasmid carrying the gene *UP2* under the control of its native promoter or, as control, the *gfp* under the constitutive promoter P_LAC_, were transferred to diverse *V. chagasii* strains. Tenfold dilutions of a phage were spotted on susceptible strain (black and red squares). Rows represent sequenced *Vibrio* strains ordered according to the Maximum Likelihood persistent genome phylogeny of *V. chagasii* (n=46) Columns represent phages (n=48) ordered by VIRIDIC clustering dendrogram. Change in susceptibility between UP2 and GFP strain strains are indicated by a red square. **C-** The susceptibility to UP2 of six phages that belong to the same VIRIDIC family than F115 was tested using their original host, i.e. the host used to isolate these phages. To this aim fold change of phage titer (as determined in A) was compared between GFP vs UP2 carrying host.

We hypothesized that PICMI_115_ immunity is mediated at least in part by a new defense system, localized between *prim* and *fis* genes and encoded by the gene *UP2*. To test this hypothesis, we cloned *UP2* under the control of its native promoter in a plasmid and transferred it through conjugation to 46 other *V. chagasii* strains that are susceptible to at least one phage^22^. As a control, the same plasmid expressing GFP was transferred to the strains. Among the 90 possible host and phage combinations that led to the production of phage progeny, eight combinations, involving eight different phages, were affected by UP2 (Fig. 5B). Six out of the eight phages affected by UP2 (Fig. 5C) belong to the same family, as defined by Virus Intergenomic Distance Calculator (VIRIDIC^34^) with pairwise identities ranging from 55 to 70%. Within this family (Fig. S17A), only the helper phage Φ115 was not affected by UP2 (Fig. S17B). *V. chagasii* strain V511 was susceptible to a member of this VIRIDIC family, Φ511 (Fig. 5C) and, to a minor extent, to genetically more diverse phages Φ168 and Φ177 (Fig. S17B). We found that UP2 influences the production of all three phages infecting the strain V511 (Fig. 5B and Fig. S17B). UP2 anti-viral activity seemed, however, dependent on the V511 genetic background, as it was not observed for the combination involving phages Φ177 and Φ168 and other hosts, including the “original host” that was used to isolate the phage (Fig. 5B). It is noteworthy that the effect of the complete PICMI_115_ element on the infection of V511 by Φ511 was much more pronounced than UP2 alone, suggesting epistatic effects within this genetic element. In the strain carrying UP2, 60 min after the addition of Φ511, the phage titer in the culture did not change (Fig. 5C, fold change 100) in contrast to the GFP control (fold change 102). In the strain carrying the full PICMI_115_ the phage titer decreased by ∼103 (Fig. 5A). We conclude that PICMI protects the bacterial host from non-helper phage. This protection relies at least in part on a novel UP2 defense system, whose activity is dependent on the host background.

## DISCUSSION

Here, we report a new family of phage satellites that is packaged as concatemers in viral particles. This results in a packaged molecule of DNA with a size similar to that of the helper phage. As a result, PICMI does not require gene(s) involved in re-shaping the capsid size. The PICMI family is among the smallest of phage satellites, with PICMI_115_ being the smallest such element with demonstrated activity. At the other end of the spectrum, cf-PICI^12^ produce their own capsids dedicated to the exclusive packaging of their genome. Their minimalist gene repertoire seems dedicated to genes for excision and integration, DNA replication, and anti- viral defenses toward competitors of the helper phage. The small size of the PICMI implies a high dependency on the helper phage for activation, packaging, and release of the particles in the bacterial lysate. In our model system of PICMI_115_, vibrio V115 and phage Φ115, this dependency is not accompanied by any significant cost for the helper phage production. This finding fits with previous results for cf-PICI, in which costs for the helper phages were also insignificant^12^.

The discovery of the PICMI family, and more specifically the mechanistic characterization of the PICMI_115_ life cycle, validates the previous hypothesis that capsid size reduction is not a common strategy for the marine satellites^7^. Furthermore, PICMI lacks identifiable packaging genes, suggesting that it does not affect the composition of the viral particle, beyond packaging it with its own DNA. What are the advantages of packaging multiple copies of the satellite within native full-sized helper phage capsids? First, non-remodeled capsids might guarantee optimal interactions with the helper phage tail. Second, it diminishes the number of functions that must be encoded by the PICMI genome. Third, having multiple copies of the satellite injected by the viral-like particle into the cell could increase the expression of satellite genes that are necessary for its integration (gene dosage), thus increasing the frequency of integration after transduction. Finally, multiple copies of the extrachromosomal element could potentially reduce the efficiency of host defense. As larger satellite size results in lower numbers of copies packaged, this finding also suggests a potential tradeoff between the acquisition of accessory genes and the efficiency of transduction. This could explain why the genes present in the *prim*- *fis* hotspot region are restricted in number and subject to high turnover. A higher efficiency of transduction by polyploidization underscores a feature that makes PICMI unique. Indeed, PLE’s transduction is severely reduced when packaged into ICP1-size capsids as concatemer or 6-7 PLE genomes relative to small remodeled capsids^9^. Future work will be needed to determine how the PICMI element is integrated or maintained as a single copy in the genome as we never detected more than one PICMI-like element in *Vibrionaceae* genomes.

PICMI induction requires the infection by its helper phage. Early after infection, the regulon encoding *alpA, UP1,* and *prim* is activated. The role of *UP1* is unknown, but it is noteworthy that *UP1* orthologs were found in 12 out of 35 subfamilies of PICMI and distributed in diverse *Vibrio* species (*V. cholerae, V. fluvialis, V. vulnificus* and *V. chagasii*) (Fig. S14). While *UP1* is not essential for PICMI activation and spread, its maintenance at 100% frequency in these subfamilies suggests it encodes for a trait that is under strong selection. *Prim,* which is necessary for efficient replication of the satellite encodes a putative RNA polymerase that synthesizes short fragments of RNA, which are then used as primers by the DNA polymerase. The *prim* gene is also found in the vast majority of known satellites, in accordance with a core and essential function of this protein in the lifestyle of satellites^13^. AlpA appears as the key regulator of the switch from latency (integrated) to activation (excised) of PICMI. Indeed, its expression is necessary and sufficient to trigger the activation of the satellite in the absence of the helper phage. Our results suggest that the PICMI *int* gene is constitutively expressed and that phage-induced *alpA* is required for the formation of a functional excision complex. Hence, several relevant questions need to be further addressed, such as: I) how does the helper phage activate the early regulon? II) How do AlpA, the integrase, and probably IOLG interact to catalyze the excision of the satellite? The understanding of *alpA* induction mechanism by the helper phage Φ115 could benefit from analysis of the genomes of closely related phages that infect the V115 strain but induce little or no PICMI.

PICMI is the second family of identified satellites induced by a virulent phage. In contrast to PLE, PICMI does not alter the production of its helper phage. We showed that PICMI can confer immunity toward other virulent phages, and we identified a new defense system (UP2) encoded by the satellite. The phage range of UP2 activity appears very narrow, and many phages susceptible to this system are phylogenetically related to the helper phages. Hence, by protecting the bacteria if the phage is not a helper, PICMI is first protecting itself. This system promotes the stable coexistence of both helper phage and satellites within the bacterial populations.

Satellite, helper phage, and bacterial host interactions are highly specific. With such a narrow host range, how do the right combinations of phages, satellites, and bacterial hosts interact in the marine environment? We speculate that blooms of specific vibrio strains can dramatically increase the abundances of specific phages and satellites. When colonizing an animal host such as oyster, vibrios can reach a higher density that might favor physical contact and promote phage infection and satellite transduction. The distribution of a satellite is expected to adhere closely to the distribution of the helper phages at a small temporal and spatial scales due to their total dependency.

We recently highlighted that many phage defense genes are encoded on large genomic islands, named phage defense elements, but the mechanisms of transfer of these elements remained unexplored^23, 28^. Around 0.6% of marine viral particles (3.2 × 1026 globally) are packaged satellites^7^, and the discovery of PICMI-mediated immunity strongly suggests that phages, including virulent ones, play an important role in the mobility of the phage defense elements. A common view is that only virulent phages should be used for phage therapy to limit horizontal gene transfer. However, our data suggest that this idea should be taken with caution because PICMI_115_ was efficiently transduced by a virulent phage. Indeed, the discovery of PICMI_115_ and its helper virulent phage underscores the importance of understanding the interactions between virulent phages and the mobile genetic elements encoded by their bacterial hosts.

## AUTHOR CONTRIBUTIONS

FLR conceived the study, supervised the project and secured funding. RBC, DP, MM and FLR conducted the experiments. DG, JMS and EPCR performed the genomic analyses. RBC, DG, JMS, DP, MM, EPCR, and FLR analysed the data. FLR, RBC, JMS and EPCR wrote the manuscript.

## ACKNOWLEDGEMENTS

We thank Agnes Thierry, Céline Loot, Francois-Xavier Barre and Mélanie Blokesch for valuable suggestions. We thank Sophie Le Panse (MERIMAGE, Roscoff), Gwen Tanguy, Erwan Legeay (GENOMER, Roscoff), Karine Cahier and Yannick Labreuche for technical assistance. We thank Jenna Sternberg from Life Science Editors for help with the Manuscript.

This work was supported by fundings from the European Research Council (ERC) under the European Union’s Horizon 2020 research and innovation program (grant agreement No 884988, Advanced ERC Dynamic) to FLR, from the Agence Nationale de la Recherche (ANR- 20-CE35-0014 « RESISTE ») to EPCR and FLR. R.B.-C. acknowledges the Spanish Ministerio de Ciencia e Innovación for his FPI predoctoral contract (BES-2017-079730). EPCR lab was funded by Equipe FRM (Fondation pour la Recherche Médicale): EQU201903007835. This work used the computational and storage services (TARS cluster) provided by the IT department at Institut Pasteur, Paris.

## METHODS

### RESOURCE AVAILABILITY

#### Lead contact

Further information and requests for resources and reagents should be directed to and will be fulfilled by the Lead Contact, Frédérique Le Roux.

#### Materials availability

Strains, phages, and plasmids generated in this study are available upon request and without restrictions from the lead contact upon request.

#### Data and code availability

Accession numbers of vibrio and phage genomes isolated and sequenced in^22^ are listed in the Key Resources table. This paper does not report original code. All programs used to analyze genomes were previously reported and are freely available online (see key resources table). The MacSyFinder models used to identify PICMI are available upon request and can be used with the MacSyFinder to make novel analysis. Any additional information required to reanalyze the data reported in this paper is available from the lead contact upon request.

### EXPERIMENTAL MODEL AND SUBJECT DETAILS

#### Bacterial strains and growth conditions

Phages and bacterial strains used in this study are listed in Table S4 and S5 respectively. Strains used or established for the genetic approach are presented in Table S6. *Vibrio. chagasii* isolates were grown in marine agar (MA, Difco) or marine broth (MB) at RT with gentle agitation. *Escherichia coli* strains were grown at 37°C in Luria-Bertani (LB, Difco) agar or in LB broth with shaking (250 r.p.m.). Chloramphenicol (Cm; 5 or 25μg/ml for *V. chagasii* and *E. coli*, respectively), thymidine (0.3 mM) and diaminopimelate (0.3 mM) were added as supplements when necessary (all chemicals from Sigma-Aldrich). Induction of the P_BAD_ promoter was achieved by the addition of 0.2% L-arabinose to the growth media, and conversely, was repressed by the addition of 1% D-glucose. Conjugation between *E. coli* and vibrios were performed at 30°C as described previously^35^ with the exception that we used TSA-2 (Tryptic Soy Agar supplemented with 1.5% NaCl) instead of LB for mating and selection. Briefly, overnight cultures of donor and recipient were diluted at 1:100 in culture media without antibiotic and grown up an OD_600nm_ of 0.3. The mating was performed on TSA-dap using a donor/recipient ratio of 5/1. Counter-selection of Δ*dapA* donor was done by plating on a TSA devoid of diaminopimelic acid (DAP) and supplemented with antibiotic.

#### Phage isolation, high titer stock, and titration

New phages infecting V115 were isolated from concentrated seawater viruses sampled in summer 2021 in the same oyster farm and using the same protocol than in^22, 23^. A volume of 100 μl of an overnight (ON) culture of bacterial host and 20 μl of viruses were directly plating on a bottom agar plate (1.5% agar, in MB) and 3.5 ml molten top agar (55°C, 0.4% agar, in MB) were added to form host lawns in overlay and allow for plaque formation^36^. Plaque plugs were first eluted in 500 µl of MB for 24 hours at 4°C, 0.2-µm filtered to remove bacteria, and re-isolated twice on V115 for purification before storage at 4°C and, after supplementation of 25% glycerol at −80°C. High titer stocks (>1011 PFU/ml) were generated by confluent lysis in agar overlays^36^. To determine the titer of phage, bacterial lawns were prepared by mixing 100 µl of on overnight culture of cells with top agar and poured onto plates. Then, tenfold dilutions of phage were spotted on plate, which were incubated at RT for 24 h.

### METHOD DETAILS

#### Plasmid construction

The primers and plasmids used or established in this study are listed in Table S7 and S8 respectively. For the preparation of quantitative PCR (qPCR) standards, each amplicon was PCR amplified using the RedTaq polymerase (VWR) and cloned in the plasmid pCR2.1 using the TOPO-TA Cloning^TM^ Kit (Invitrogen).

For vibrio mutagenesis, cloning was performed using Herculase II fusion DNA polymerase (Agilent) for PCR amplification and the Gibson Assembly Master Mix (New England Biolabs, NEB) for insert-plasmid assembly, according to the manufacturer instructions. All cloning was confirmed by digesting plasmid minipreps with specific restriction enzymes and/or sequencing (Eurogentec).

#### Nucleic acid extraction, amplification, and southern blot

Prior to DNA extraction, phage suspensions (5 ml, >1011 PFU/ml) were concentrated to approximately 500 µl on centrifugal filtration devices (30 kDa Millipore Ultra Centrifugal Filter, Ultracel UFC903024) and washed with 1/100 MB to decrease salt concentration. Alternatively, phages were concentrated using PEG 8000 1X and NaCl 1M, incubated ON at 4°C, centrifuged 30 min at 4500 rpm, and the pellet was resuspended in 500 µl SM buffer (NaCl 100 mM, MgSO4.7H20 8 mM, Tris-Cl 50 mM). The concentrated phages were next treated for 30 min at 37°C with 10 µl of DNAse (Promega) and 2.5 µl of RNAse (Macherey-Nagel) at 1000 unit and 3.5 mg/ml, respectively. The nucleases were inactivated by adding EDTA (20 mM, pH 8). DNA extraction encompassed a first step of protein lysis (0.02 M EDTA pH 8.0, 0.5 mg/ml proteinase K, 0.5% sodium dodecyl sulfate) for 30 min incubation at 55°C, a phenol chloroform extraction, and an ethanol precipitation. Bacterial DNA was extracted using the Wizard Genomic DNA Purification Kit (Promega).

RNA was extracted with TRIzol^TM^ Reagent (Sigma-Aldrich) and High Pure RNA Isolation Kit (Roche), treated by TURBO DNAse (Ambion) and reverse transcripted using the Transcriptor First Strain cDNA Synthesis Kit (Roche).

Classical PCRs were performed using the RedTaq (WVR) and amplicons were visualized by SYBR Green stained (Sigma) agarose gel electrophoresis (1 to 2% agarose). qPCR and qRT-PCR was performed using LightCycler 480 SYBR Green I Master (Roche). The thermal cycling conditions were 95°C for 10 min, followed by 40 cycles of 95°C for 10 s, 60°C for 20 s and 72°C for 25 s, then 1 cycle of 95°C for 5 s, 65°C for 1 min and 95°C for 15 s. Standard curves were constructed using serial dilutions of plasmid, leading to the number of DNA copies per 20 ng of DNA. Number of copies for phage, empty integration site, and circularized PICMI_115_ were normalized by the number of copies of vibrio (*gyrB*) per sample. For the qRT-PCR, the resulting copies number were normalized on *gyrA* for each sample. To determine fold change, samples collected at different time post-infection were compared to the sample before adding phages.

For Southern blot, DNA samples were run on 0.7% agarose gel at 100V for one hour. Then, the DNA was transferred to Nylon membranes (Hybond-N+; Amersham Life Science) using standard methods. DNA was detected using a DIG-labelled probe (Digoxigenin-11-dUTP alkali-labile), anti-DIG antibody (Anti-Digoxigenin-AP Fab fragments) and Chemiluminescent detection with CSPD following the instructions of the kit (all products and kits from ROCHE).

#### Construction of HiC libraries, sequencing, and analysis

1 ml of different mix of high titer stocks (>1011 PFU/ml) of phages (mix1: 1 ml of Φ115, mix2: 1 ml of Φ191, mix3: 500 µl of Φ115 + 500 µl of Φ191) were fixed in a 5 ml Eppendorf tube by adding formaldehyde (Sigma-Aldrich, ref -F4775, Formalin 35-36.5% plus methanol 15%) to a final concentration of 3% and incubated at RT for 1 hour under gentle agitation. The reaction was stopped by adding glycine (stock = 2.5 M) to a final concentration 0.125 M and incubated at RT for 20 min under gentle agitation. Fixed particles were then centrifuged at 16,000 x g for 20 minutes at 4°C. Supernatant was discarded, resuspended in 1 ml of PBS 1X, and recentrifuged at 16,000 x g for 20 minutes at 4°C. The supernatant was again discarded carefully and the pellet was resuspended in 45 µl of Tris 10 mM pH 7.5. The HiC libraries were then constructed using the ARIMA Kit (Arima Genome-Wide HiC+ Kit). HiC genomic libraries were then processed for sequencing as previously described^37^ and were sequenced on Nextseq550 apparatus (2 x 35 bp). Contact maps were generated using Hicstuff^38^ (bowtie2 - very sensitive local mode – mapping quality of 30) and a reference FastA files containing the 3 phage genomes. Contact maps were then binned at 1kb resolution, balanced, and displayed using Hicstuff.

#### Electron microscopy

Following concentration on centrifugal filtration devices (Millipore, amicon Ultra centrifugal filter, Ultracel 30K, UFC903024), 20 µl of the phage concentrate were adsorbed for 10 min to a formvar film on a carbon-coated 300 mesh copper grid (FF-300 Cu formvar square mesh Cu, delta microscopy). The adsorbed samples were negatively contrasted with 2% Uranyl acetate (EMS, Hatfield, PA, USA). Imaging was performed using a Jeol JEM-1400 Transmission Electron Microscope equipped with an Orious Gatan camera at the platform MERIMAGE (Station Biologique, Roscoff, France).

#### Vibrio mutagenesis

PICMI labelling (PICMI_115_-Cm^R^) was performed by cloning the 500bp end of UP2 gene in the suicide plasmid pSW23T^39^. To inactivate UP2 or *prim* gene and at the same time label the PICMI derivatives (Δprim and ΔUP2 -Cm^R^), a 500bp internal region of the gene was cloned in the suicide plasmid pSW23T. After conjugative transfer, selection of the plasmid-borne drug marker (Cm^R^) resulted from integration of pSW23T in the target region by a single crossing-over. The integration of the suicide plasmid was verified by PCR.

Gene deletion was performed by cloning 500bp fragments flanking the gene in the pSW7848T suicide plasmid^40^. This pSW23T derivative vector encodes the *ccdB* toxin gene under the control of an arabinose-inducible and glucose-repressible promoter, P_BAD35_. Selection of the plasmid-borne drug marker on Cm and glucose resulted from integration of pSW7848T in the genome. The second recombination leading to pSW7848T elimination was selected on arabinose-containing media. Mutants were screened by PCR using external primers.

For the complementation experiments, the genes necessary for PICMI_115_ excision, *int/IOLG* and *alpA,* or a *gfp* control were cloned under the control of the conditional P_BAD_ promoter in a P15A-*ori*-based replicative vector. The plasmids were transferred by conjugation into the mutants. Strains were grown to mid-exponential phase in the presence of 0.2% arabinose (activation of P_BAD_) and then infected with Φ115pure for 30 min. To explore its anti-phage activity, UP2 under its native promoter was cloned in a pMRB plasmid^41^, and the same plasmid expressing the GFP was used as control.

We were not able to delete the complete PICMI115 by allelic exchange using the pSW7848T suicide plasmid. As an alternative, we cloned the *alpA* gene of the *V. chagasii* or *V. aestuarianus* under the control of P_BAD_ promoter in a P15A-*ori*-based replicative vector (Spec^R^), assuming that the expression of *alpA* in trans is sufficient to induce PICMI_115_ excision even in the absence of phage (Fig. 2C) and the *alpA* from the two *Vibrio* species are interchangeable for PICMI_115_ induction (Table S9). The plasmid was transferred by conjugation to V115. The Spec^R^ conjugant was grown overnight in MB with Spec and arabinose, serially diluted and plated on TSA-2 (TSB-2 with agar). A total of 48 colonies were screened by PCR to identify V115 derivatives that lack the PICMI (ΔPICMI). We obtained clones with the mutation (8/48) only when using *alpA* from *V. aestuarianus*.

#### PICMI induction

The vibrio strain was grown to mid-exponential phase in Marine broth (OD=0.3) and infected under static condition with the phage at a multiplicity of infection (MOI) of 10 otherwise indicated. At each time point, an aliquot of the culture was centrifuged, the supernatant was filtered at 0.2 μm and the titer of phages was determined by drop spotting serial dilutions of the supernatant on the host lawn. To determine fold change, the titer of phage in the lysate was compared to the same amount of phage added in the culture media without bacteria. Total RNA and/or DNA was extracted from the bacterial pellet.

#### Adsorption estimation

Phage adsorption experiments were performed as previously described^42^. Phages were mixed with exponentially growing cells (OD 0.3; 107 CFU/ml) at a MOI of 0.01 and incubated at RT without agitation. At different time points, 250 µl of the culture was transferred in a 1.5 ml tube containing 50 µl of chloroform and centrifuged at 14,000 rpm for 5 min. The supernatant was 10-fold serially diluted and drop spotted onto a fresh lawn of a sensitive host to quantify the remaining free phage particles. In this assay, a drop in the number of infectious particles at 15 or 30 min indicated bacteriophage adsorption.

#### PICMI transduction

The number of PICMI particles were quantified using the transduction tittering assay. Briefly, lysates were produced by infecting V115 derivatives carrying PICMI_115_-Cm^R^ and derivatives by Φ115pure. A 1:100 dilution (in fresh MB broth) of an overnight recipient strain was grown until an OD_600_ of 0.3 was reached. Then, 100 ml of the recipient culture was dispatched in a 96 well plate, infected by addition of 10 µl of PICMI lysate serial dilutions prepared with MB for 1H at RT. The different mixtures of culture-PICMI-Cm^R^ were plated out on TSA-2 plates containing chloramphenicol. LBA plates were incubated at RT for 24 h, and the number of colonies formed (transduction particles present in the lysate) were counted and represented as the colony forming units (CFU/ml). PCRs were performed to confirm the integration of PICMI at the end of the *fis* gene.

#### *In silico* prediction and analysis of PICMI-like element

The PICMI-like elements were searched using two datasets: 1) the bacterial division of GenBank release v243 (5/26/2021) that contains 24,243 complete genomes, including 456 genomes of the *Vibrionaceae* family and 2) the NCBI Assembly database (2/16/2023) with 19,185 *Vibrionaceae* genomes available but with variable assembly quality. SatelliteFinder (v0.9.1)^13^ was used on both datasets with a dedicated PICMI model defined by four mandatory genes encoding: the integrase (PF00589.25, PF00239.24, PF07508.16), AlpA (PF05930.15, PF12728.10), the primase (DUF3987 with PF13148.9 and DUF5906 with PF19263.2) and the Fis regulator (PF02954.22). The resulting elements were then filtered by excluding those with *int, alpA,* and *fis* localized in different contigs, those predicted to belong to other families (PICI, cf-PICI, P4 and PLE), and those integrated into a gene showing a lower identity with *fis.* The genomic region starting with the *fis* gene and ending with the direct repeat upstream the *int* gene was extracted, aligned with FAMSA (v1.6.2)^43^, and each PICMI subfamily was defined by a pairwise nucleic identity ≥90% (Table S3 and Fig. S14).

The PICMI genes were clustered in families using mmseqs2 (v14.7e284) reciprocal best-hit^44^ with 20% identity and 50% coverage thresholds (Table S3 and Fig. S14). Phage defense systems were annotated using Defense-Finder (v1.0.8 and models v1.1.0)^33^ and phage structural genes were annotated using PhANNs (v1.0.0)^45^ with a threshold score ≥7.

The functional annotation of genes used multiple approaches, i.e. tblastn similarity searches on GenBank, InterPro domain prediction^46^, and ProtNLM annotation [https://www.uniprot.org/help/ProtNLM]. The 3D protein structures predicted by ColabFold (v1.5.2-patch)^30^ were compared to publicly available protein structures using the Foldseek search server (v5)^47^.

Comparative genomics were performed using PanACoTA workflow (v1.4.0)^48^. Persistent genes were defined as present in single copy in at least 90% genomes with a minimum of 30% protein identity. Protein sequences of each family were first aligned and concatenated. Phylogenetic reconstruction was done using iqtree2 (v2.0.3)^49^ with 1000 bootstraps and GTR model. Genome plots were generated using dedicated python scripts based on the ‘DNA Features Viewer’ library (https://github.com/Edinburgh-Genome-Foundry/DnaFeaturesViewer).

We clustered phages using VIRIDIC (v1.1, default parameters)^34^. Intergenomic similarities were identified using BLASTN pairwise comparisons. Viruses’ assignment into genera (**≥**70% similarities) and species (**≥**95% similarities) ranks follows the International Committee on Taxonomy of Viruses (ICTV) genome identity thresholds. We used PHANOTATE v1.5.0^50^ for the syntaxic annotation of phage Φ115 and, after removing restriction on gene size, analysis of large non-coding regions within PICMIs.

### Nanopore genome assembly and analysis

The Nanopore sequencing library was prepared using Native Barcoding genomic DNA (EXP-NBD104) and Ligation Sequencing Kit 1D (SQK-LSK109) and sequenced using MinION flow cell R9.4.1 at the platform GENOMER (Station Biologique, Roscoff, France).

Demultiplexing and base calling of raw nanopore sequencing data (Table S1) was performed using Guppy software (v6.1.1, --flowcell FLO-MIN106 --kit SQK-LSK109). The base called sequences were used as input for genome assembly performed using FLYE (v2.9)^51^ RAVEN (v1.4.0)^52^ and NECAT (v0.0.1)^53^ with default parameters.

The comparisons of Illumina and Nanopore assemblies of both Φ115 and PICMI_115_ were performed using pairwise FAMSA alignment (v1.6.2)^43^. FLYE was selected for further analysis because it showed the highest similarity with previous Illumina sequencing. Nanopore PICMI reads were analyzed using a dedicated script based on FAMSA alignment to precisely determine read start and end on an artificial 13 copies concatemer reference.

