## Supplemental Figures for "Phage inducible chromosomal minimalist island (PICMI), a family of satellites of marine virulent phages"

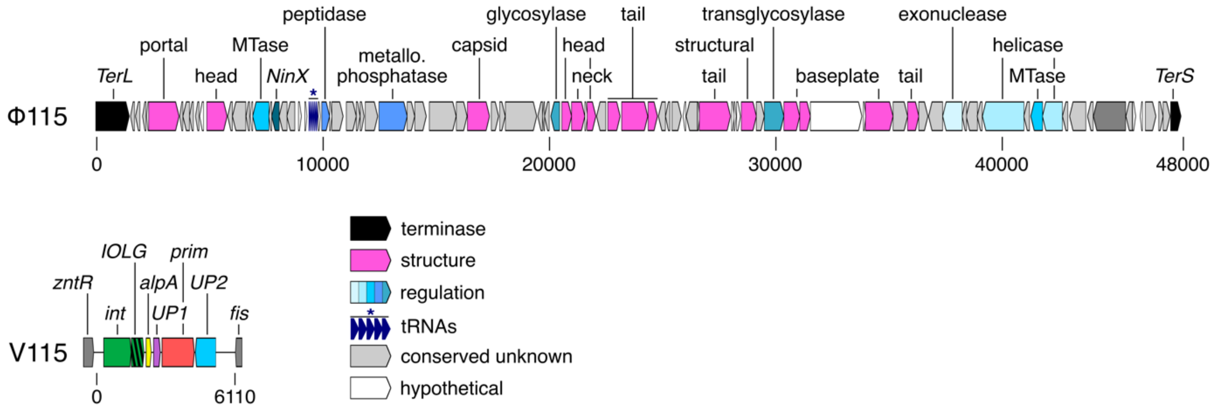


**Figure S1**. Genome assembly and annotation of the phage 115_E_34-1 (Φ115) revealed two contigs of 47,851 and 6,110 bp respectively. The larger contig corresponds to the complete genome of the phage, ordered here from the large (TerL) to the small (TerS) terminase (locus tags VP115E341_P0001 to 0093). The small contig were also found in the genome of the host used to isolate and propagated this phage, *V. chagasii* 34_P_115 (V115). This region is integrated in the bacterial genome at the end of the *fis* regulator gene (locus tag VCHA34P115_150113) and flanked by two direct repeats of 17 bp (aatacggcatgaactaa). Among the six genes identified in this region, *int* encodes for an integrase, *alpA* for a putative regulator and *prim*, a putative primase.

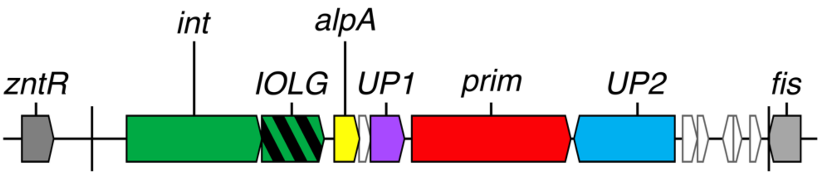


**Figure S2.** Syntactic annotation of the satellite using phanotate without size threshold suggest six additional putative genes (in white) that were considered as false open reading frame (ORF).

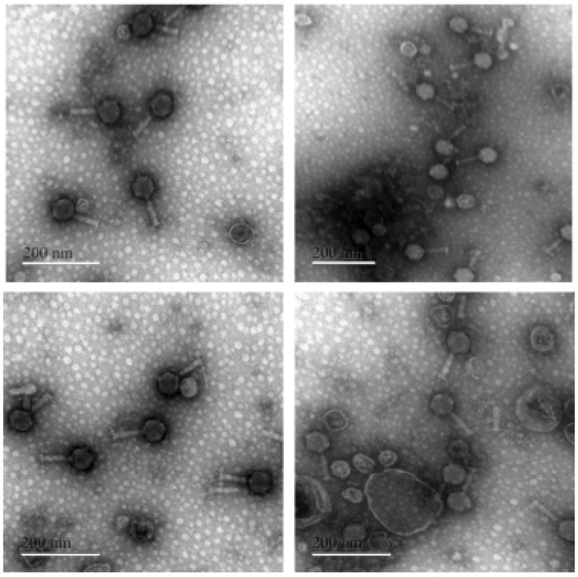


**Figure S3.** Transmission electron micrographs of Φ115 myovirus, produced using *V. chagasii* V115 as host. Images are representatives of the observation of dozens of particles that did not revealed heterogeneity in capsid size. Images were collected using a Jeol JEM-1400 TEM Microscope. Scale bars are 200 nm.

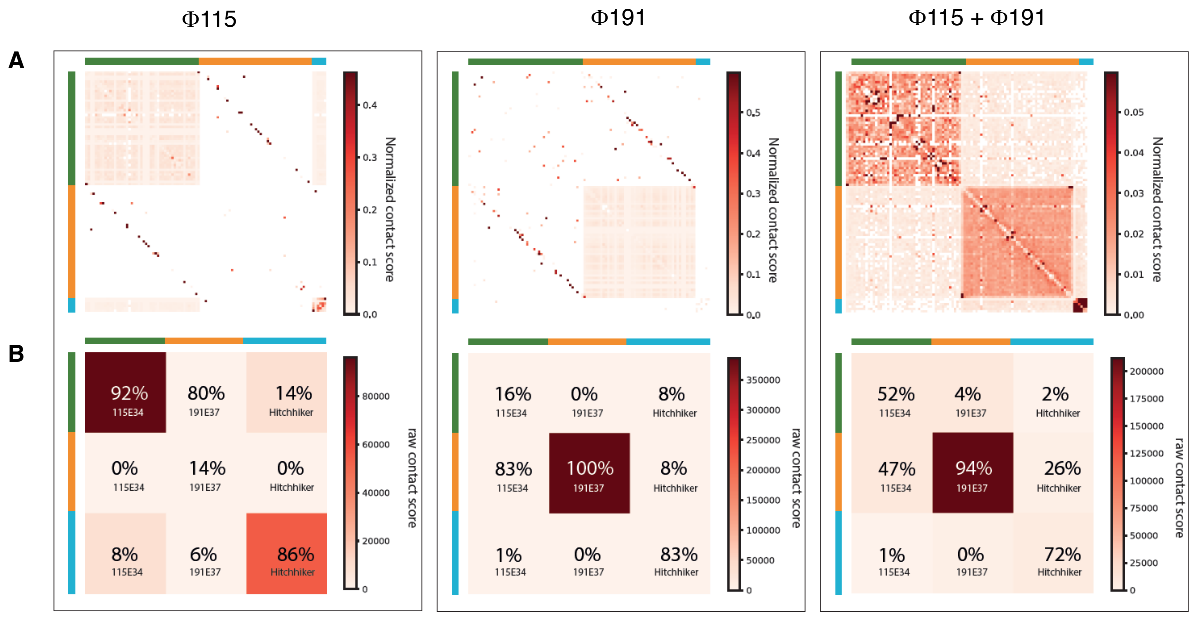


**Figure S4.** HiC experiment demonstrating that the genome of the phage Φ115 and its satellite are located in distinct viral particles.

**A-** Chromosomal contact maps of phage Φ115, phage Φ191 another myovirus with similar genome size and a single genome, or a mixture 1:1 of both phages. The color code represents normalized contacts between DNA regions, ranging from low (white) to high (red) frequencies (1pixel = 1kb, color scale: vmax = 99% of maximal value). The genome sequence of the helper phage Φ115 and its satellite is indicated in green and blue respectively. The genome sequence of the phage Φ191 is in orange. The result supports the absence of physical contact between the phage Φ115 and the satellite DNAs. The observed contacts result from background. All three genomes show a signal in the upper right and lower left corners indicating circular molecules or a circular permutation phenomenon.

**B-** Contact map representing the raw contact signal for the same three experiments. Proportions of contact in each category are indicated in percentage relatively to total number of contacts for each genomic entity.

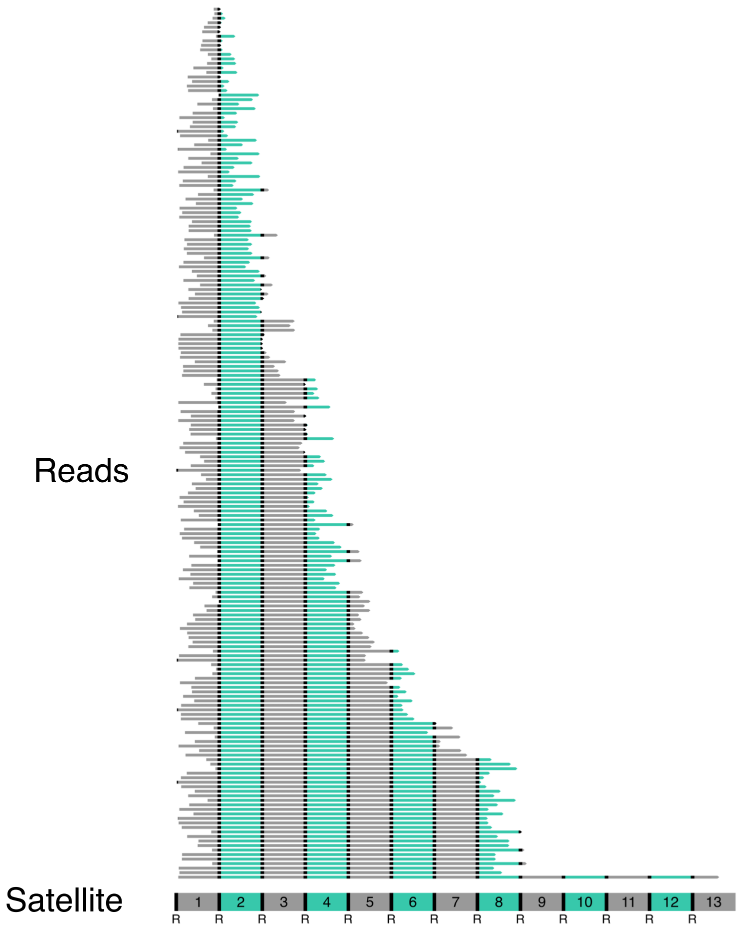


**Figure S5.** Nanopore reads mapped to $PICMI_{115}$ concatemer structure. Nanopore sequencing of Φ115 revealed that a fraction of the viral particles contained a concatemer of 8 copies of the 6.1 kbp satellite leading to ~49 kbp size. All obtained reads were aligned with FAMSA to an artificial $PICMI_{115}$ concatemer of 13 copies, corresponding to the longest read length. Each copy of $PICMI_{115}$ concatemer is represented by alternating grey/green, and overlapping sequences correspond to the direct repeats (AATACGGCATGAACTAA, indicated as "R").

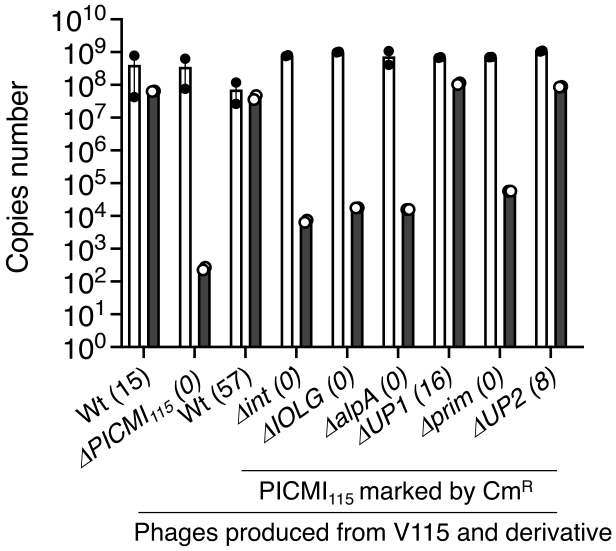


**Figure S6.** Estimation of the percentage of viral particles that contain phage satellites instead of phage DNA with a population. The number of copies of phage Φ115 (white bar) or concatemer of the satellite (grey bar) were determined by qPCR using DNAs extracted from a high titer of phages produced in V115 wild type (wt) and a derivative lacking the entire satellite ($\Delta$PICMI$_{115}$). For transduction experiments, a chloramphenicol resistance marker ($Cm^R$) was introduced within the satellite of V115 and derivatives lacking one of the six satellite genes (e.g. Δ*int*). Bar charts show the mean +/- SEM. from two technical replicates experiments (individual dots). On the x axis, number in brackets indicates the estimate percentage of the Φ115-like particles hitchhiked by the satellite.

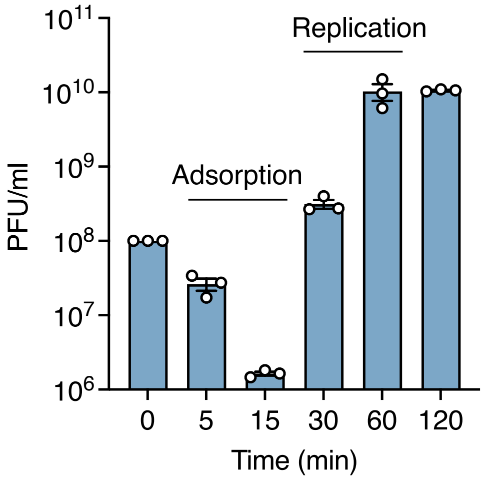


**Figure S7.** Estimation of the infection dynamic by Φ115. *V. chagasii* strain V115 was grown to mid-exponential phase in Marine broth (OD=0.3) and infected with pure Φ115 at a multiplicity of infection (MOI) of 10. Aliquot of the culture was centrifuged at the indicated times, the supernatant was filtered at 0.2 µm, and the titer of phages was determined by drop spotting serial dilutions of the supernatant on the host lawn. Bar charts show the mean +/- SEM from three independent experiments (individual dots).

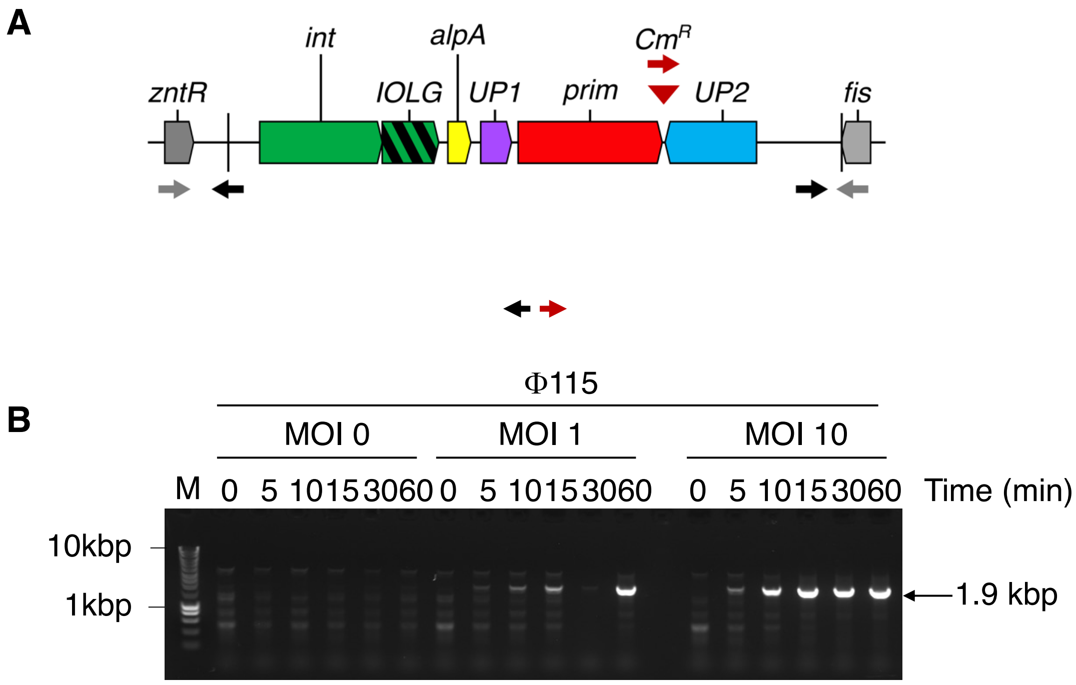


**Figure S8. PICMI$_{115}$ is induced following Φ115 infection**.

**A-**Schematic representation of PICMI$_{115}$ integrated between the *fis* and *zntR* genes in the genome of *V. chagasii* strain V115. For transduction assays, this element was marked by a chloramphenicol resistance cassette (brown triangle). Arrows depict Forward and Reverse primers used to detect the circularized and concatemeric form of PICMI$_{115}$ (1972 bp).

**B-** A phage Φ115 stock was produced using V115 as host and thus contains ~15% of particles containing PICMI$_{115}$. The V115 host strains was grown to mid-exponential phase in Marine broth (OD=0.3) and infected with Φ115 at a multiplicity of infection (MOI) of 0, 1 and 10. At the indicated times (in minutes) cells were pelleted, total DNA was extracted and used as template for PCR to detect the circularized and concatemeric form of PICMI$_{115}$.

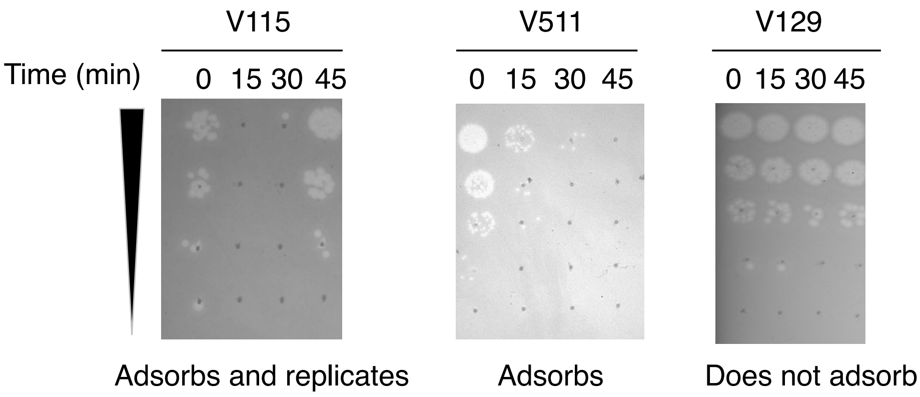


**Figure S9**. Estimation of the adsorption of Φ115 on three *V. chagasii* strains. After allowing a fixed concentration of phages (MOI 0.01) to adsorb to each *V. chagasii* strain for the indicated time, free phages that remained unattached were serially diluted and plated with the original host, V115. In this assay, a drop in the number of infectious particles indicates phage adsorption. Adsorption was complete after 15 or 30 minutes using respectively V115 and V511 as host and was not observed up to 45 minutes using V129 (negative control). After 45 minutes, production of phages was observed only when using V115.

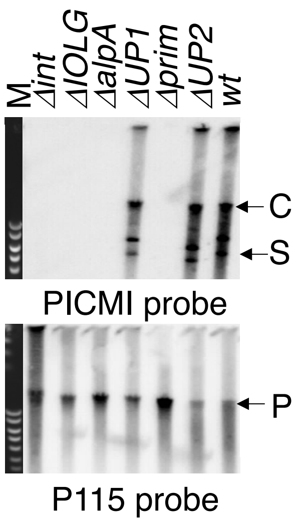


**Figure S10.** Genes necessary for $PICMI_{115}$ activation. The strain V115 and derivatives lacking one of the six genes from $PICMI_{115}$ were infected by the phage Φ115pure for 30 minutes. DNA was extracted from bacteria separated on a 0.7% agarose gel and Southern blotted with $PICMI_{115}$ or Φ115 probes. M: molecular marker (Smart ladder Eugentec). C, S and P indicate the concatemeric and single form of $PICMI_{115}$ and phage genome respectively.

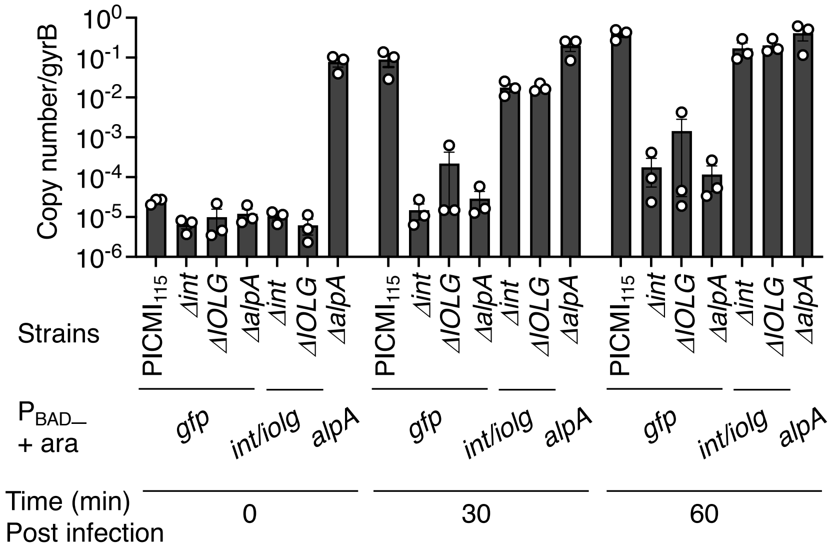


**Figure S11**. AlpA is sufficient to induce $PICMI_{115}$ activation. The $PICMI_{115}$ genes, *int/IOLG* and *alpA* or, as control, *gfp* were cloned under the control of the conditional $P_{BAD}$ promoter and the plasmids were transferred by conjugation into the respective mutants. Strains were grown to mid-exponential phase in the presence of 0.2% arabinose (activation of $P_{BAD}$) and then infected with Φ115pure for the indicated time. The circular from of $PICMI_{115}$ was detected by qPCR and normalized on vibrio (*gyrB)* copies number. Bar charts show the mean +/- SEM from three independent experiments (individual dots). The expression of *alpA* is sufficient to induce $\Delta PICMI_{115}$ activation in the absence of phage (upper panel, time 0 min).

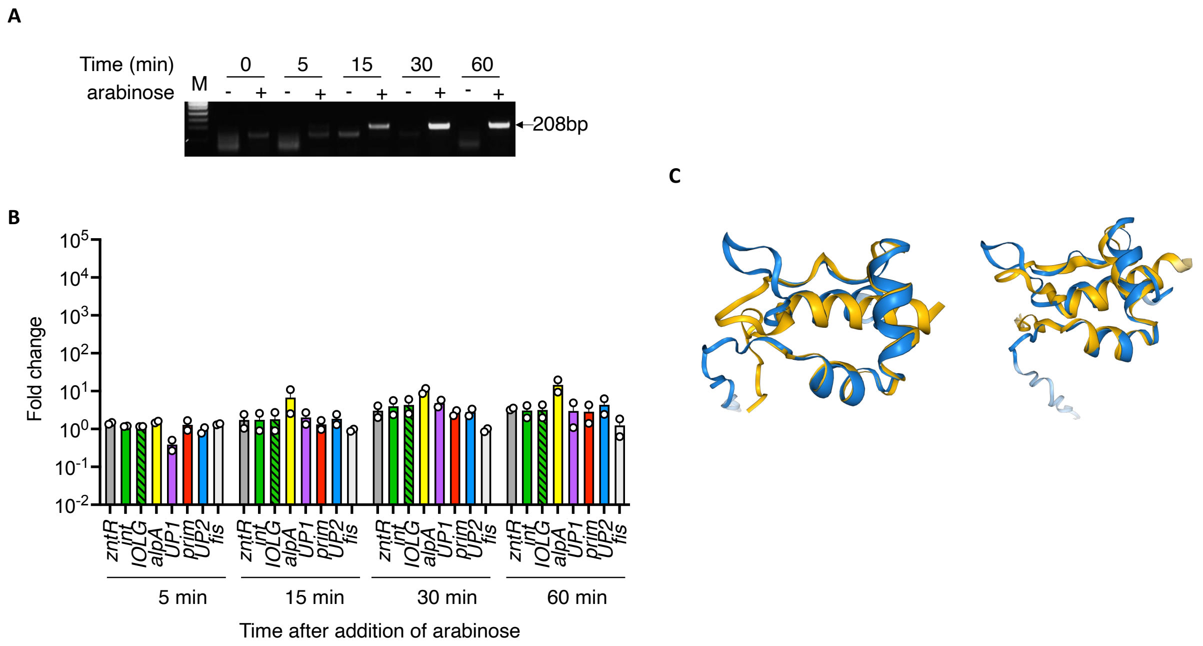


**Figure S12.** AlpA is involved in the formation of the excision complex rather than as a transcriptional regulator of $PICMI_{115}$ genes. The gene *alpA* cloned in a plasmid under the control of the conditional $P_{BAD}$ promoter was transferred by conjugation into the ΔalpA mutant. The strain was grown to mid-exponential phase in Marine broth (OD=0.3) and arabinose (0.2%) was added (+) or not (-) as indicated. At the indicated time, total DNA and RNA were extracted.

**A-** PCR and gel stained to detect the circularized and concatemeric form of $PICMI_{115}$.

**B-** qRT-PCR to detect the expression of each of the six genes from $PICM_{I115}$, as well as the two flanking genes *fis*, *zntR,* and the house keeping gene *gyrA*. The resulting copies number were normalized on *gyrA*. Bar charts show the mean +/- SEM from two independent experiments (individual dots). With the exception of *alpA* expressed in trans from the plasmid, we did not find significant difference between presence and absence of arabinose, indicating transcriptional activation of PICMI genes by alpA is unlikely.

**C-** Structure superposition of AlpA and TorI response regulator (left) and Xis excisionase (right). AlpA from PICMI (in blue) structure is predicted using ColabFold and search against AlphaFold/Swiss-Prot v2 database. One of the top 5 hits, TorI response regulator inhibitor for tor operon (Q1R904) from *Escherichia coli* strain UTI89 (TM-score=0.6585) (left) or Xis excisionase (P15482) from *Streptomyces ambofaciens* (TM-score=0.74838) (right) are superimposed in yellow.

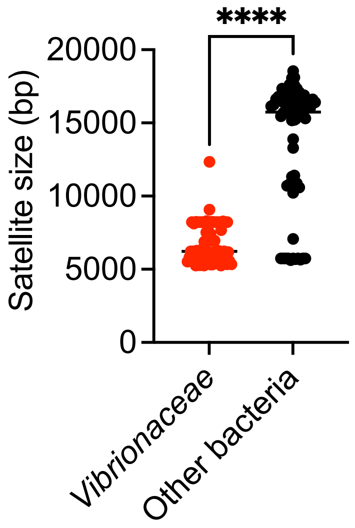


**Figure S13.** Comparison of the size of the 135 putative satellites, identified by the colocalization of *int, alpA, prim,* and *fis* genes, in the Genbank bacterial genome (v243), including 67 elements found in *Vibrionaceae* genomes.

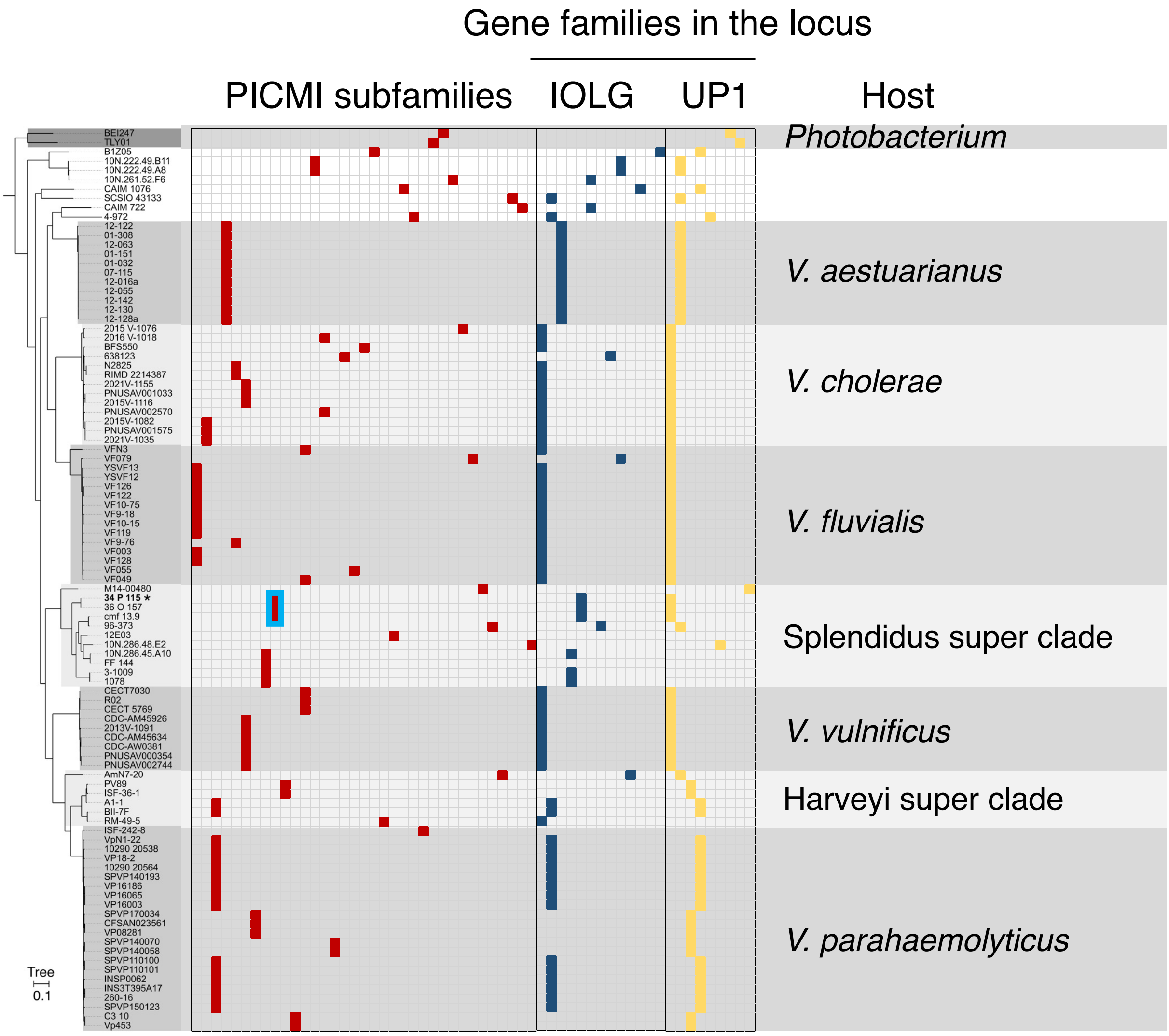


**Figure S14.** Genes frequently found across the PICMI subfamilies in the *Vibrionaceae.* A persistent core phylogenetic tree was constructed using PanACoTA (90% for the persistent genome resulting to 1026 families and 30% for minimum percentage of identity) with iqtree2 (1000 bootstrap and GTR model). Among the 35 PICMI subfamilies (35 columns, brown squares), 21 carry an integrase overlapping gene (IOLG), and four elements carry a gene contiguous to the *int* gene (ICG). The *IOLG* or *ICG* were grouped in 13 distinct gene families (13 columns, dark blue squares). Together with the core genes *alpA* and *prim, UP1* is part of the early regulon activated by the helper phage. Homologs of $PICMI_{115}$ UP1were found in 12 PICMIs. Some single genes from eight distinct families, encoding for unknown function, were also present between *alpA* and *prim* in other PICMIs (total 9 gene families in the locus where we found UP1 in $PICMI_{115}$). The subfamily corresponding to $PICMI_{115}$ is framed in light blue.

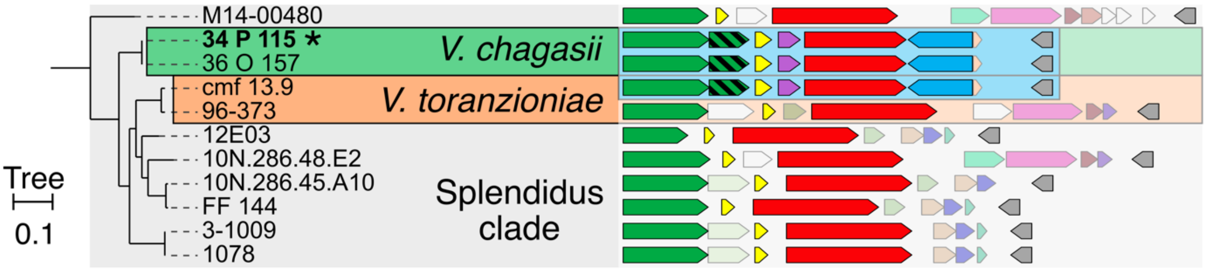


**Figure S15.** The PICMI$_{115}$ subfamily was detected in two strains of *V. chagasii* (34_P_115 and 35_O_157 named V115 and V157 for simplicity) and in a *V. toranzoniae* strain (cmf 13.9) within the Splendidus clade (zoom in the phylogenetic tree from Fig. 4 and S14). Another *V. toranzoniae* (96-373) carries a different PICMI subfamily.

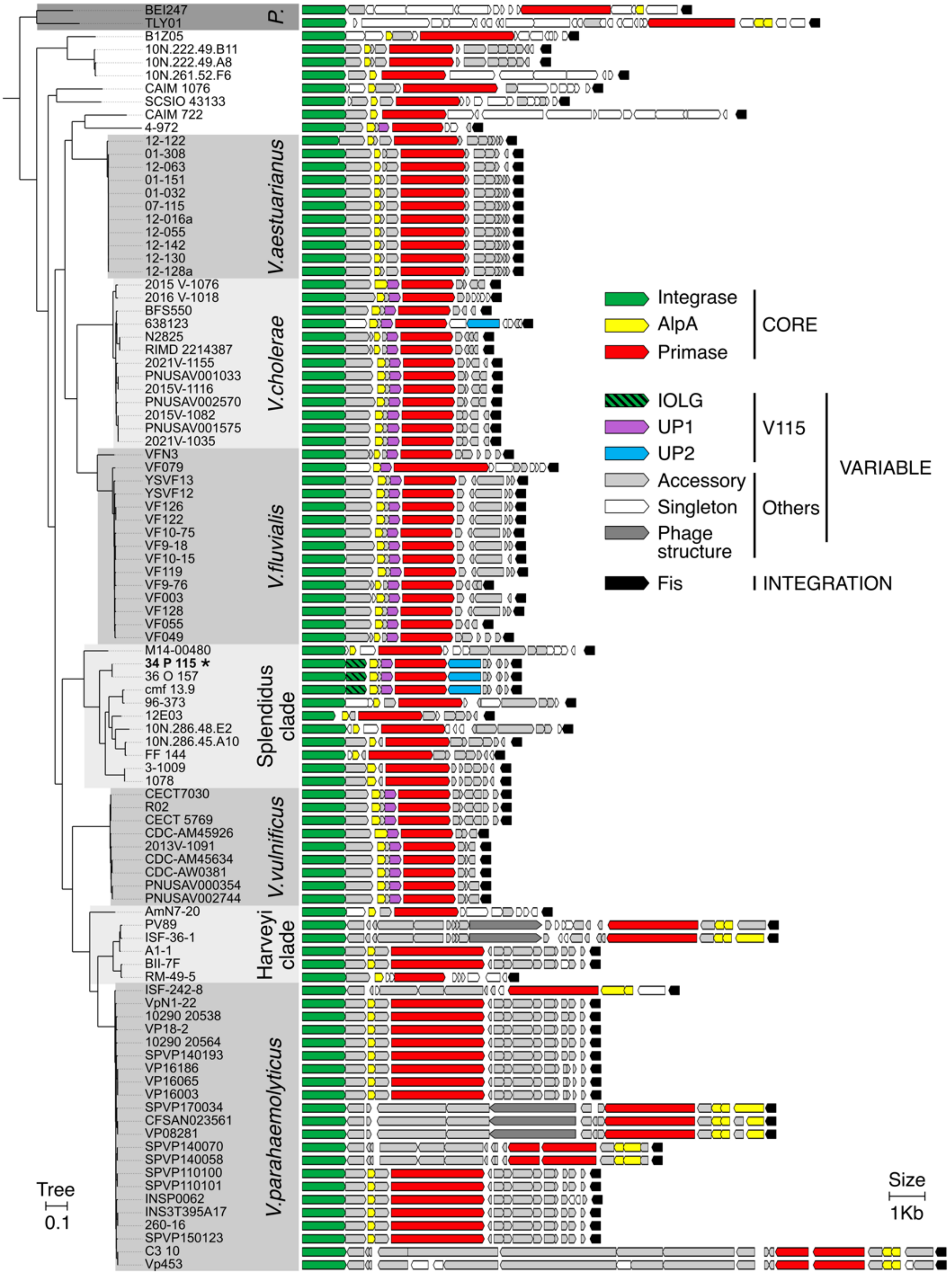


**Figure S16.** Phylogenetic persistent core tree and genomic representation of the 97 PICMI elements found in *Vibrionaceae* (GenBank 01-27-23 containing 19189 organisms). The persistent core tree was constructed using PanACoTA (90% for the persistent genome resulting to 1026 families and 30% for minimum percentage of identity) with iqtree2 (1000 bootstrap and GTR model). Genus, super clades, or species names are indicated in the grey boxes. "P." corresponds to *Photobacterium* genus, Harveyi, Splendidus are super clades encompassing several *Vibrio* species. The $PICMI_{115}$ element is pinpointed by bold strain name (34_P_115, V115) and by an asterisk. PICMI-like elements were searched using SatelliteFinder and the identified genomic regions were reannotated using phanotate without length filter. This led to the syntaxic annotation of several small ORFs (<50 amino acids), including in $PICMI_{115}$, that were considered as fragmented and/or pseudogenes. The region from the integrase to the *fis* gene was plotted using the DnaFeaturesViewer python library. Solid colors indicate core $PICMI_{115}$ genes. Grey colors indicate accessory and singleton PICMI-like genes defined using reciprocal best-hit with 20% identity for 50% coverage.

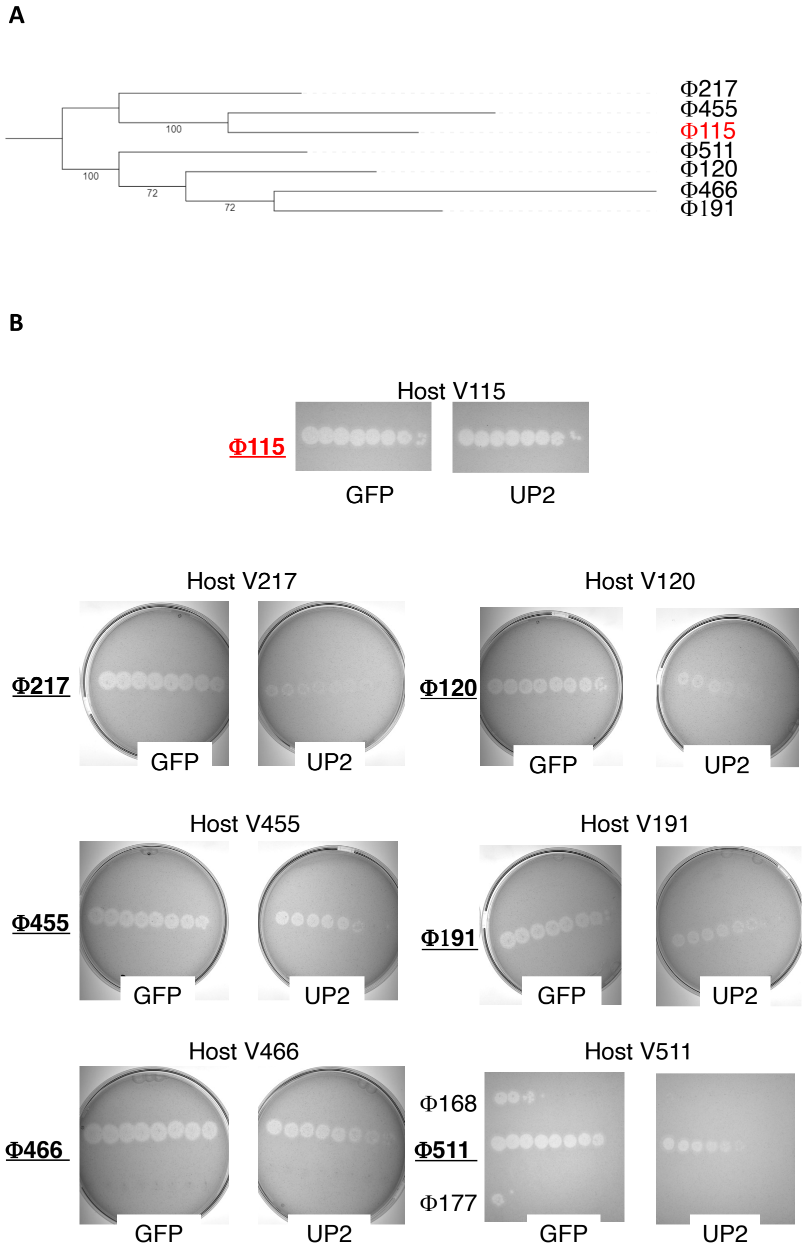


**Figure S17.** Changes in susceptibility to phage killing observed for UP2 expressing *V. chagasii*.
**A-** Core phylogenetic tree for phages from the VIRIDIC family (>50% identities) including the helper phage Φ115 (red). The phylogenic tree is based on core proteins (25% identities and 80% coverage).
**B-** Tenfold dilutions of the indicated phages (bold underlined, from the same VIRIDIC family) were spotted on the respective host carrying a plasmid with the gene *UP2* under the control of its native promoter or, as control, the *gfp* under the constitutive promoter $P_{LAC}$. Images are representative of two independent experiments.

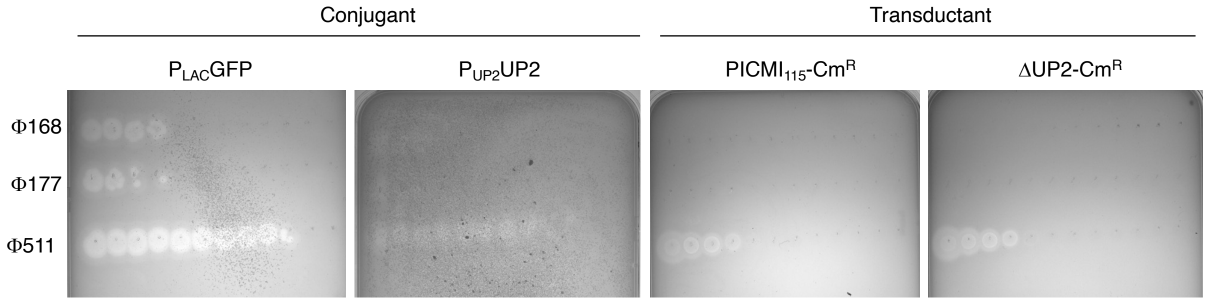


**Figure S18**

Changes in susceptibility to phage killing observed for UP2 expressing *V. chagasii* (conjugant) or carrying $PICMI_{115}$ (transductant). Tenfold dilutions of the indicated phages were spotted on V511 conjugants with plasmid containing the *gfp* under a constitutive promoter ($P_{LAC}$GFP) or the gene *UP2* under the control of its native promoter ($P_{UP2}$UP2) and transductants carrying the full satellite ($PICMI_{115}$-$Cm^R$) or a derivative with an inactivated UP2 ($\Delta$UP2-$Cm^R$). Images are representative of two independent experiments.
